# Smooth Muscle Cell-Specific TGFβ2 Protects Against Thoracic Aortic Aneurysm and Dissection in Mice

**DOI:** 10.1101/2025.10.01.679917

**Authors:** Mengistu G. Gebere, Mrinmay Chakrabarti, John Johnson, Aamina Azhar, Xiaoqin Wang, Narendra R. Vyavahare, Mohamad Azhar

**Affiliations:** Department of Cell Biology and Anatomy, School of Medicine, University of South Carolina, Columbia, SC, USA; Department of Internal Medicine, Yale Cardiovascular Research Center, Section of Cardiovascular Medicine, Yale University, New Haven, CT, USA; Department of Biology, Furman University, Greenville, SC, USA; Biomedical Engineering, Clemson University, Clemson, SC, USA

**Author notes:** Correspondence: Mohamad Azhar, PhD, 6439 Garners Ferry Road, Columbia SC 29208.

**Keywords:** transforming growth factor beta, smooth muscle, aneurysm, aorta, mice, Loeys- Dietz syndrome

## Abstract

**Objective:** Thoracic aortic aneurysm and dissection (TAAD) are major complications of Loeys-Dietz syndrome caused by heterozygous *TGFB2* mutations. While *Tgfb2* knockout mice die at birth and adult heterozygotes develop late, non-dissecting or non-rupturing aneurysms, the role of vascular smooth muscle cell (SMC)–derived TGFβ2 in postnatal aortic homeostasis and disease remains undefined.

**Approach and Results:** We generated tamoxifen-inducible, SMC-specific *Tgfb2* conditional knockout mice (*Tgfb2*cKO) by crossing *Tgfb2*^flox^ alleles with *Myh11CreER*^T2^ and *ROSA*^mT/mG^ lineage reporter mice. *Tgfb2* deletion was induced at 4 weeks of age. *Tgfb2*cKO mice developed rapidly progressive aneurysms involving both ascending and descending aortas, with intramural dissection and/or rupture at the proximal descending aorta. Lineage tracing confirmed loss of *Tgfb2*-deficient SMCs during disease progression. Histological and morphometric analyses revealed elastic fiber fragmentation, SMC loss and de-differentiation, medial thickening, adventitial fibrosis, and accumulation of collagen and proteoglycans. Molecular profiling demonstrated reduced expression of SMC contractile genes (*Acta2, Myh11*), increased fibrillar collagen (*Col1a1*) expression, early suppression of SMAD2/3 phosphorylation and increased non-canonical TGFβ signaling via p38 and pERK1/2 MAPK pathways.

**Conclusions:** These findings demonstrate that TGFβ2 derived from vascular SMCs is essential for postnatal aortic wall homeostasis by preserving SMC differentiation, maintaining extracellular matrix integrity, and supporting and preserving a proper balance of both canonical and non-canonical TGFβ signaling. Loss of SMC-specific *Tgfb2* precipitates medial degeneration, aneurysm formation, dissection, and rupture, providing direct mechanistic insight into *TGFB2*-associated aortopathy and establishing a robust novel genetic mouse model for evaluating targeted therapies in TAAD.

**Highlights:**

- Postnatal, SMC-specific *Tgfb2* deletion in mice caused rapidly progressive thoracic aortic aneurysms, dissections, and fatal rupture.
- Loss of *Tgfb2* disrupts SMC contractile phenotype and ECM homeostasis, leading to medial degeneration, elastin fragmentation, and abnormal collagen/proteoglycan accumulation.
- Canonical TGFβ–SMAD signaling is suppressed, while MAPK pathways are activated, indicating ligand-specific signaling imbalance.
- Findings highlight TGFβ2 as a central regulator of postnatal aortic homeostasis and suggest that targeted ligand-specific therapeutic strategies may better preserve aortic wall stability.

**Significance:** This study identifies smooth muscle cell–derived TGFβ2 as a critical, nonredundant regulator of postnatal aortic wall integrity, linking its loss to thoracic aortic aneurysm, dissection, and rupture, and highlighting TGFβ2 ligand-specific signaling as a targeted therapeutic target.

**Graphical Abstract:** Smooth muscle cell–derived TGFβ2 maintains postnatal aortic wall homeostasis by preserving contractile gene expression, elastin architecture, and balanced ECM remodeling. Conditional deletion of *Tgfb2* in SMCs shifts signaling from canonical SMAD2/3 to MAPK pathways, leading to medial degeneration, progressive aneurysm, dissection, and rupture—highlighting TGFβ2 as a nonredundant, ligand-specific regulator and potential therapeutic target in thoracic aortopathy.

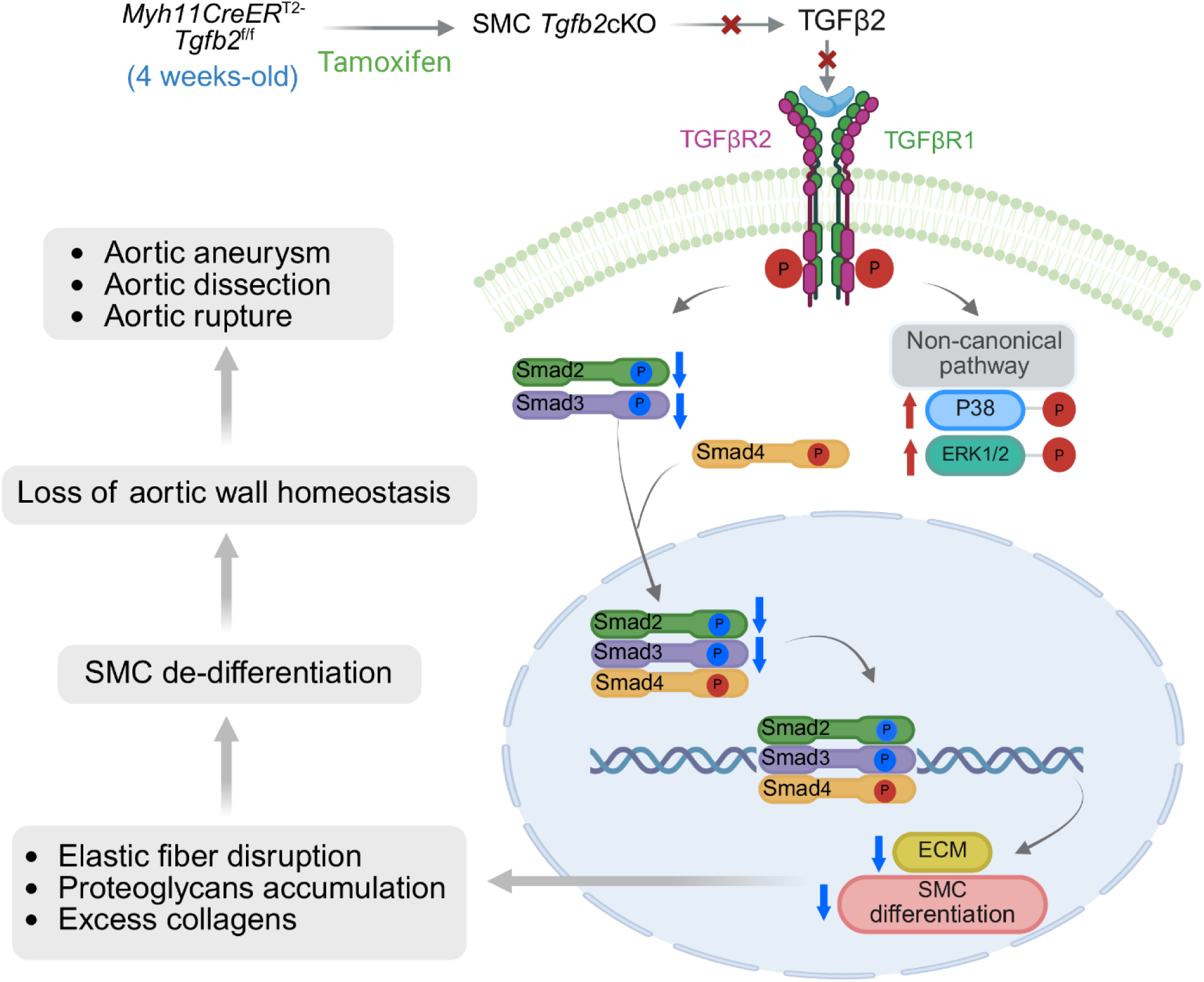

## Introduction

Thoracic aortic aneurysms and dissections (TAAD) are progressive disorders of the aortic wall characterized by degeneration of medial elastic fibers and loss of structural integrity^1^. TAAD is responsible for more than 10,000 deaths annually in the United States^2^. A thoracic aortic aneurysm (TAA) represents a permanent and irreversible dilation of the aorta, whereas aortic dissection involves disruption of medial elastic layers with hemorrhage into and along the wall, often culminating in rupture. Acute dissection and rupture are the most lethal complications, with an estimated incidence of 2.53 per 100,000 patient-years, and dissections may arise both in the presence and absence of aneurysmal dilation ^1, 3, 4^. Heritable disease is subdivided into syndromic and nonsyndromic forms, depending on whether extracardiac manifestations are present.

Approximately 20% of TAA cases are familial ^5, 6^. Mutations in smooth muscle contractile genes such as *ACTA2, MYH11, MYLK*, and *PRKG1* account for many nonsyndromic cases, while syndromic forms are associated with connective tissue disorders including Marfan syndrome (MFS), Loeys-Dietz syndrome (LDS), vascular Ehlers-Danlos syndrome (vEDS), and osteogenesis imperfecta^7^. Acquired risk factors such as aging, smoking, trauma, and cardiovascular disease also contribute to sporadic aneurysms^1^. Syndromic and nonsyndromic TAAD share several fundamental pathogenic mechanisms despite differences in clinical presentation. Both forms are characterized by medial degeneration of the aortic wall, including smooth muscle cell (SMC) loss, elastic fiber fragmentation, and accumulation of proteoglycans^3^. Genetic mutations affecting components of the extracellular matrix (ECM), smooth muscle contractile proteins, and the transforming growth factor beta (TGFβ) signaling pathway are implicated in both syndromic (e.g., Marfan syndrome, Loeys-Dietz syndrome) and nonsyndromic cases, indicating overlapping molecular pathways^8^. In both types, abnormal SMC function and dysregulation of ECM homeostasis contribute to progressive weakening of the aortic wall. This structural vulnerability, combined with mechanical stress from blood flow, predisposes the aorta to dilation and eventual dissection^9–11^. Additionally, both forms can exhibit incomplete penetrance and variable expressivity, underscoring the complex interplay between genetic and environmental factors in TAAD pathogenesis. Current guidelines recommend prophylactic surgery when the ascending aorta exceeds 5.5 cm or grows more than 0.5 cm/year, but outcomes suggest earlier intervention may improve survival – particularly in patients of LDS^12, 13^. Pharmacological therapy to prevent TAAD remains unavailable, highlighting the need for deeper insight into the molecular mechanisms - particularly in specific genetic context - underlying disease initiation and progression^2, 14^.

TGFβ ligands are multifunctional cytokines that regulate proliferation, migration, differentiation, apoptosis, adhesion, and ECM synthesis ^15–17^. The three isoforms (TGFβ1, TGFβ2, TGFβ3) which are product of three different genes show distinct yet overlapping expression in the media and adventitia of the developing aorta ^18^. *In vivo* studies demonstrate nonredundant roles for each ligand in development and tissue homeostasis, but their specific contributions to aortic maturation and adult homeostasis remain poorly understood^19^. Following activation, which is tightly controlled by ECM proteins such as A Disintegrin And Metalloproteinase with Thrombospondin Motifs 6 (ADAMTS6), fibrillins, and latent TGFβ-binding proteins (LTBP1- 4), all ligands signal through a common TGFβRII/TGFβRI (ALK5) receptor complex, sometimes with type III coreceptors such as betaglycan (i.e., TGFβR3) or endoglin^20–22^. TGFβ ligands interact with their receptors to activate the transcription factors SMAD2 and SMAD3, which become phosphorylated in the cytoplasm and subsequently accumulate in the nucleus, where they bind to enhancer or promoter elements within regulatory regions of TGFβ target genes to either induce or repress transcription ^23^. Even though TGFβ signal through common receptors, it is not clear how disruption of TGFβ ligands leads to unique and overlapping phenotypes in mice and humans^24–26^.

Given that none of TGFβ ligands deficient mice survive to adulthood, it is unknown whether homozygous deletion of individual TGFβ ligands in vascular SMCs is sufficient to cause thoracic aortic aneurysm. *TGFB2* and *TGFB3* mutations are a major cause of LDS, which shares aneurysm and dissection phenotypes with MFS and Shprintzen-Goldberg syndrome (SGS) ^27, 28^ ^26, 29, 30^. The loss of function mutations in patients with *TGFB2* mutations (OMIM#614816, LDS4) leads to decreased *TGFB2* levels and dysfunctional aortic elastin deposition and medial wall degeneration resulting in TAAD. Pathogenic variants in *TGFB2* has also been reported in the early age onset of sporadic thoracic aortic dissection (ESTAD) individuals without a family history of thoracic aortic disease or syndromic features ^5, 6^. *Tgfb2*-null mice die perinatally, while heterozygotes develop aortic dilation only after 8 months without dissection or rupture^27, 31^. The role of TGFβ2 specifically in vascular SMCs during postnatal aortic homeostasis has not been investigated. To address this gap, we deleted *Tgfb2* in SMCs of young mice and analyzed histological, molecular, and biochemical alterations during the onset and progression of thoracic aortic aneurysm and dissection. These findings identify TGFβ2 as a critical regulator of postnatal aortic wall integrity and suggest that therapeutic approaches aimed at preserving smooth muscle cell function and restoring extracellular matrix balance may improve outcomes for patients with *TGFB2*-related thoracic aortic disease.

## Main Methods

Expanded methods are provided in the Supplemental Material. All reagents, resources, and instrumentation used are provided in extended Tables (1-3)

**Table 1.**
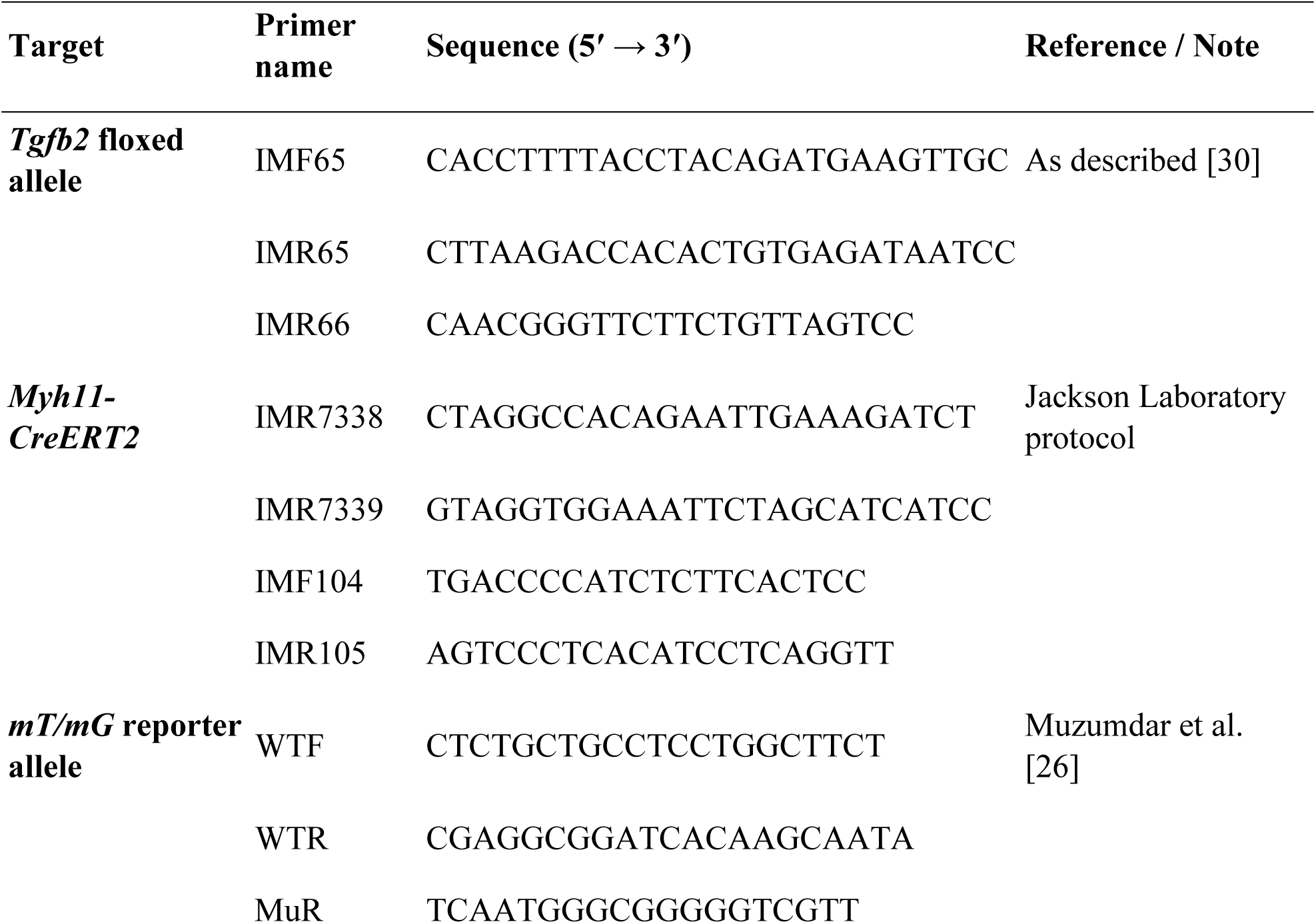
Primers used for genotyping.

**Table 2.**
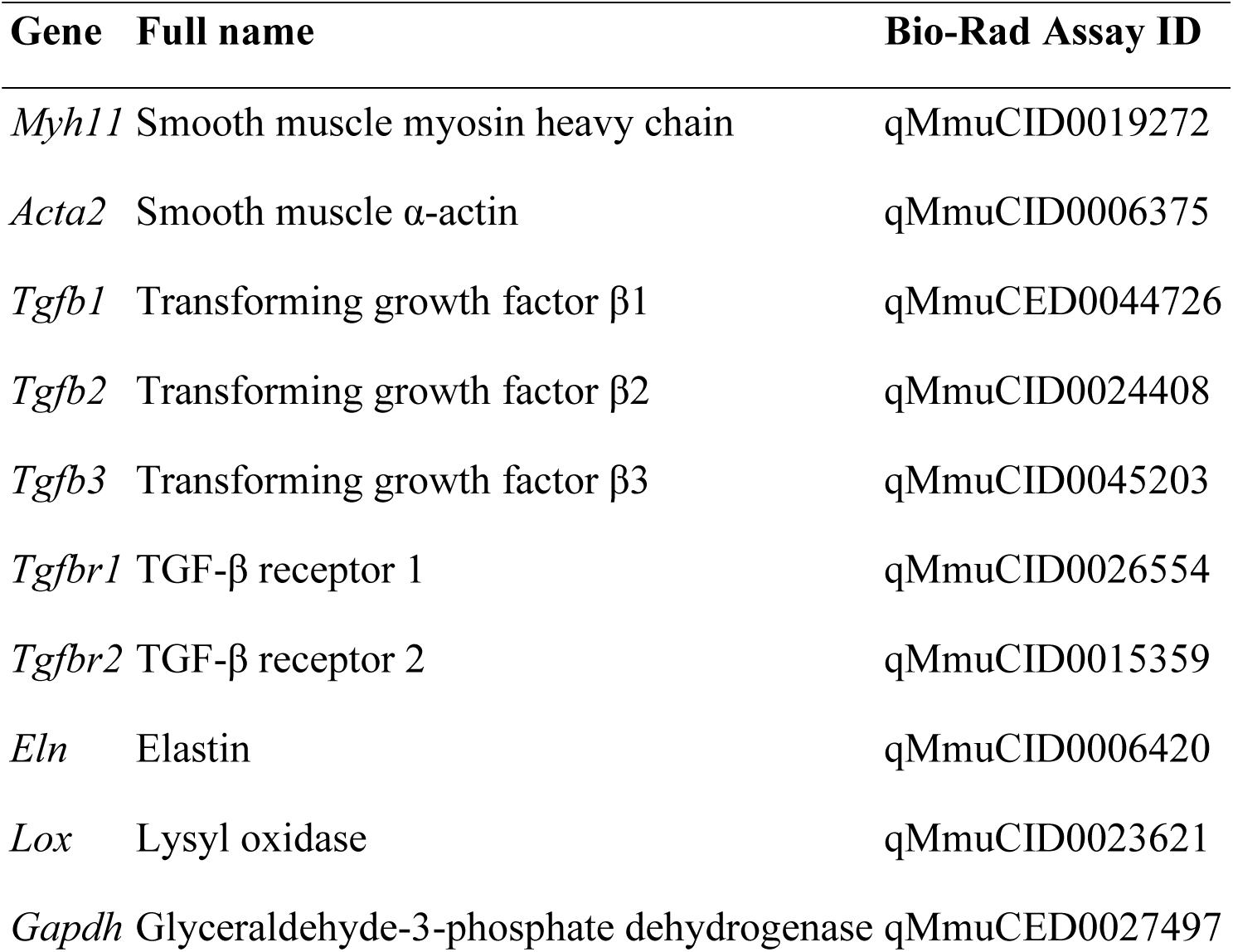
Primers used for quantitative real-time PCR (Bio-Rad PrimePCR assays)

| **Gene** | **Full name** | **Bio-Rad Assay ID** |
| --- | --- | --- |
| *Myh11* | Smooth muscle myosin heavy chain | qMmuCID0019272 |
| *Acta2* | Smooth muscle α-actin | qMmuCID0006375 |
| *Tgfb1* | Transforming growth factor β1 | qMmuCED0044726 |
| *Tgfb2* | Transforming growth factor β2 | qMmuCID0024408 |
| *Tgfb3* | Transforming growth factor β3 | qMmuCID0045203 |
| *Tgfbr1* | TGF-β receptor 1 | qMmuCID0026554 |
| *Tgfbr2* | TGF-β receptor 2 | qMmuCID0015359 |
| *Eln* | Elastin | qMmuCID0006420 |
| *Lox* | Lysyl oxidase | qMmuCID0023621 |
| *Gapdh* | Glyceraldehyde-3-phosphate dehydrogenase | qMmuCED0027497 |

**Table 3.**
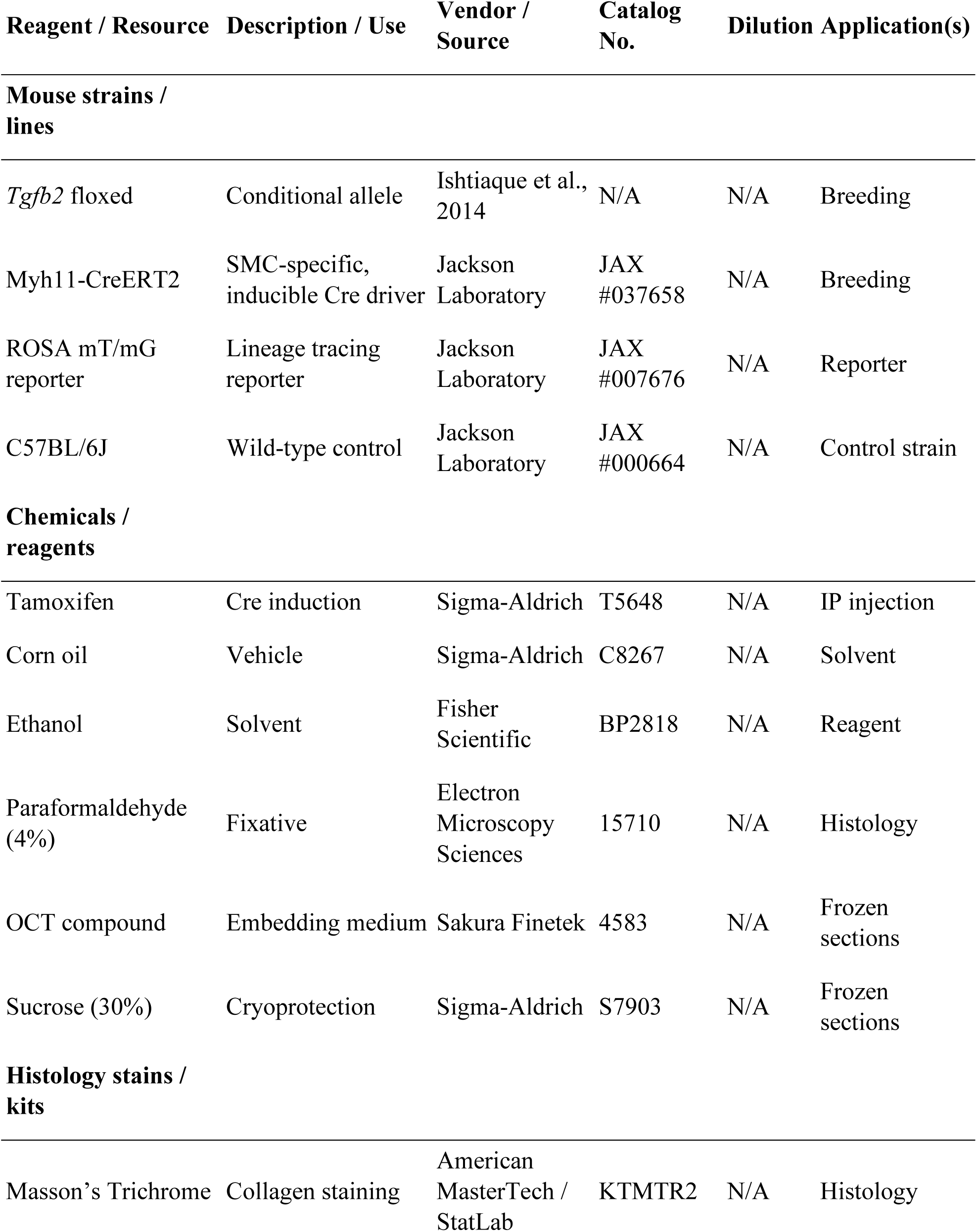
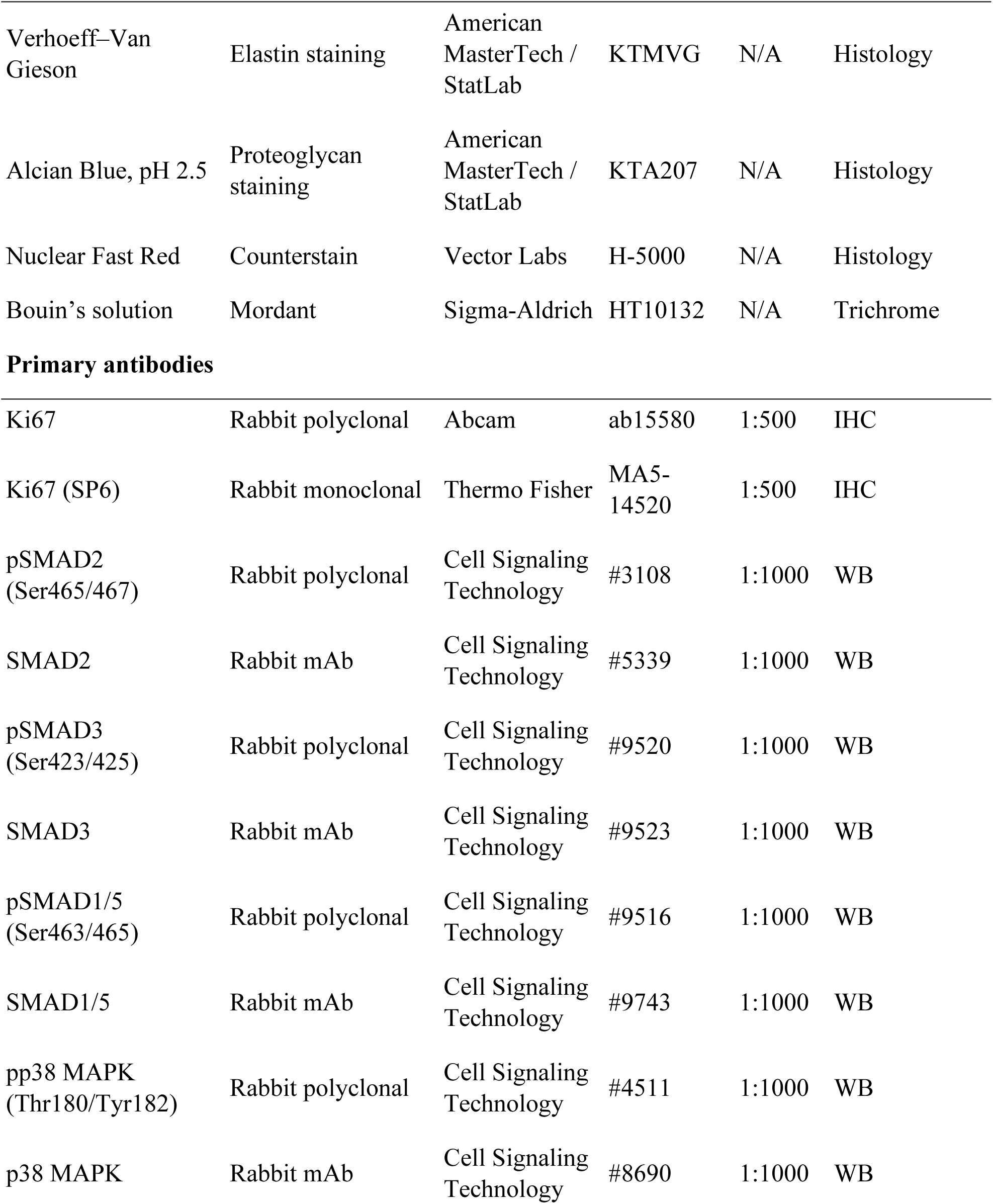
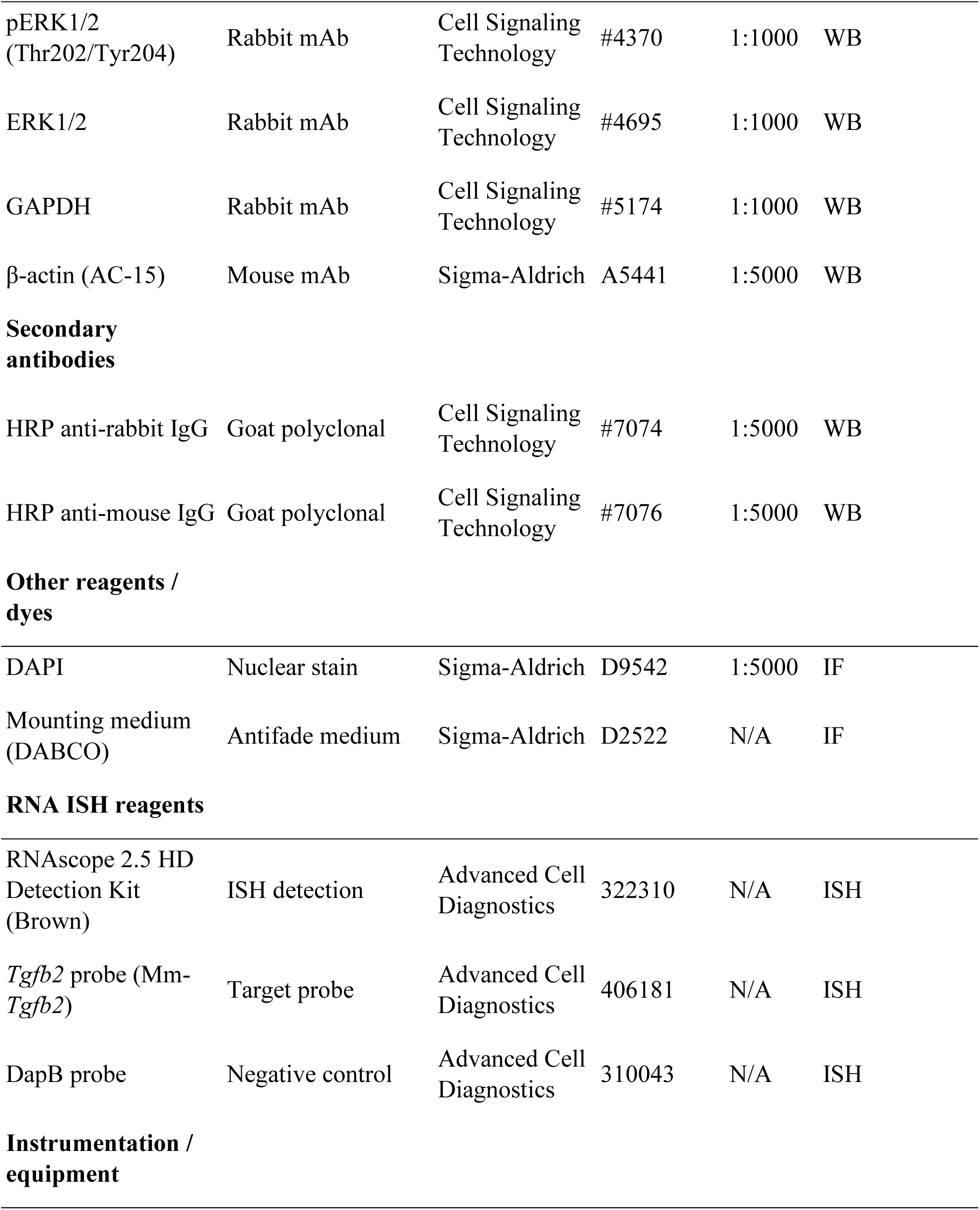
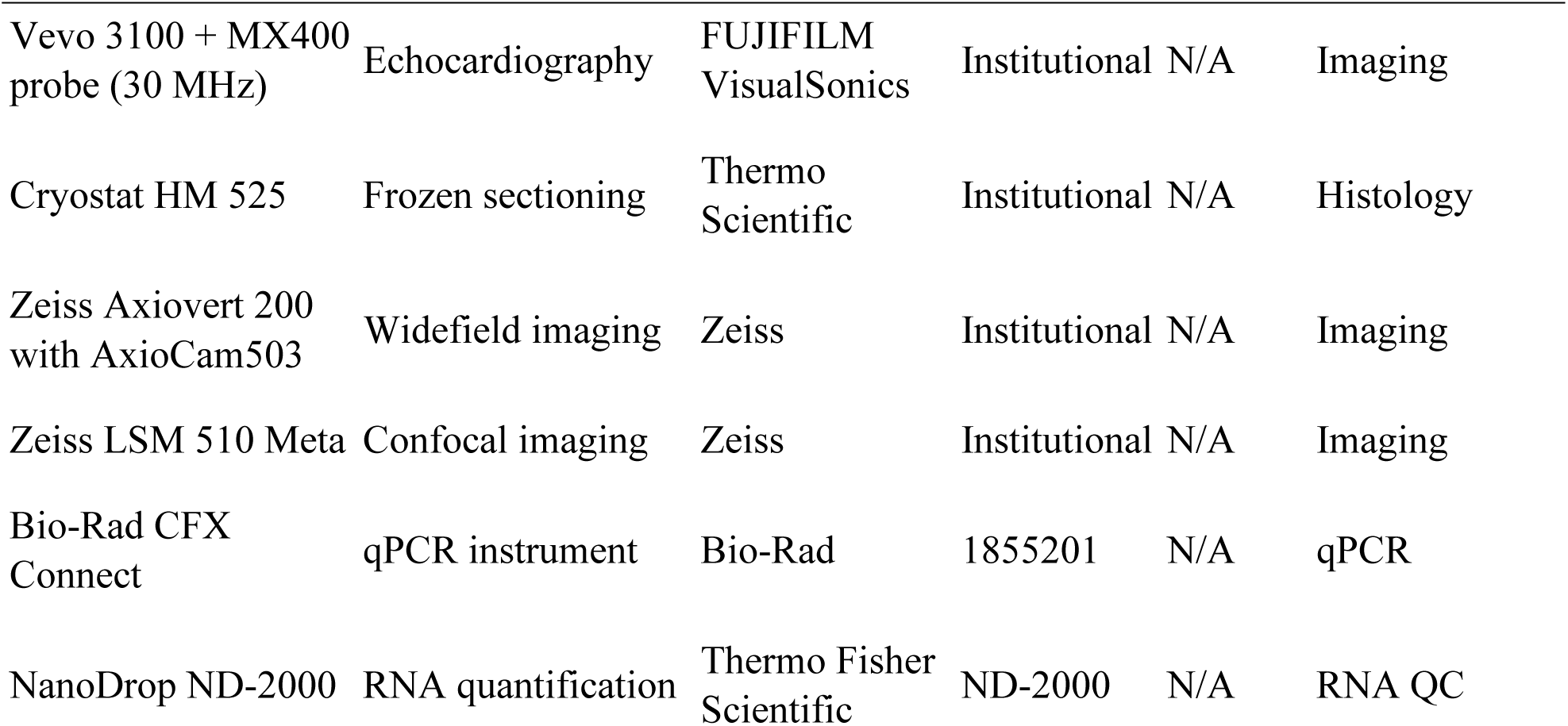
Key reagents, organisms, and resources with RRIDs.

| **Reagent / Resource** | **Description / Use** | **Vendor / Source** | **Catalog No.** | **Dilution** | **Application(s)** |
| --- | --- | --- | --- | --- | --- |
| **Mouse strains / lines** |  |  |  |  |  |
| *Tgfb2* floxed | Conditional allele | Ishtiaque et al., 2014 | N/A | N/A | Breeding |
| Myh11-CreERT2 | SMC-specific, inducible Cre driver | Jackson Laboratory | JAX #037658 | N/A | Breeding |
| ROSA mT/mG reporter | Lineage tracing reporter | Jackson Laboratory | JAX #007676 | N/A | Reporter |
| C57BL/6J | Wild-type control | Jackson Laboratory | JAX #000664 | N/A | Control strain |
| **Chemicals / reagents** |  |  |  |  |  |
| Tamoxifen | Cre induction | Sigma-Aldrich | T5648 | N/A | IP injection |
| Corn oil | Vehicle | Sigma-Aldrich | C8267 | N/A | Solvent |
| Ethanol | Solvent | Fisher Scientific | BP2818 | N/A | Reagent |
| Paraformaldehyde (4%) | Fixative | Electron Microscopy Sciences | 15710 | N/A | Histology |
| OCT compound | Embedding medium | Sakura Finetek | 4583 | N/A | Frozen sections |
| Sucrose (30%) | Cryoprotection | Sigma-Aldrich | S7903 | N/A | Frozen sections |
| **Histology stains / kits** |  |  |  |  |  |
| Masson's Trichrome | Collagen staining | American MasterTech / StatLab | KTMTR2 | N/A | Histology |

| Reagent / Resource | Description / Use | Vendor / Source | Catalog No. | Dilution | Application(s) |
| --- | --- | --- | --- | --- | --- |
| Verhoeff–Van Gieson | Elastin staining | American MasterTech / StatLab | KTMVG | N/A | Histology |
| Alcian Blue, pH 2.5 | Proteoglycan staining | American MasterTech / StatLab | KTA207 | N/A | Histology |
| Nuclear Fast Red | Counterstain | Vector Labs | H-5000 | N/A | Histology |
| Bouin’s solution | Mordant | Sigma-Aldrich | HT10132 | N/A | Trichrome |
| **Primary antibodies** |  |  |  |  |  |
| Ki67 | Rabbit polyclonal | Abcam | ab15580 | 1:500 | IHC |
| Ki67 (SP6) | Rabbit monoclonal | Thermo Fisher | MA5-14520 | 1:500 | IHC |
| pSMAD2 (Ser465/467) | Rabbit polyclonal | Cell Signaling Technology | #3108 | 1:1000 | WB |
| SMAD2 | Rabbit mAb | Cell Signaling Technology | #5339 | 1:1000 | WB |
| pSMAD3 (Ser423/425) | Rabbit polyclonal | Cell Signaling Technology | #9520 | 1:1000 | WB |
| SMAD3 | Rabbit mAb | Cell Signaling Technology | #9523 | 1:1000 | WB |
| pSMAD1/5 (Ser463/465) | Rabbit polyclonal | Cell Signaling Technology | #9516 | 1:1000 | WB |
| SMAD1/5 | Rabbit mAb | Cell Signaling Technology | #9743 | 1:1000 | WB |
| pp38 MAPK (Thr180/Tyr182) | Rabbit polyclonal | Cell Signaling Technology | #4511 | 1:1000 | WB |
| p38 MAPK | Rabbit mAb | Cell Signaling Technology | #8690 | 1:1000 | WB |
| pERK1/2<br>(Thr202/Tyr204) | Rabbit mAb | Cell Signaling Technology | #4370 | 1:1000 | WB |
| ERK1/2 | Rabbit mAb | Cell Signaling Technology | #4695 | 1:1000 | WB |
| GAPDH | Rabbit mAb | Cell Signaling Technology | #5174 | 1:1000 | WB |
| β-actin (AC-15) | Mouse mAb | Sigma-Aldrich | A5441 | 1:5000 | WB |
| **Secondary antibodies** |  |  |  |  |  |
| HRP anti-rabbit IgG | Goat polyclonal | Cell Signaling Technology | #7074 | 1:5000 | WB |
| HRP anti-mouse IgG | Goat polyclonal | Cell Signaling Technology | #7076 | 1:5000 | WB |
| **Other reagents / dyes** |  |  |  |  |  |
| DAPI | Nuclear stain | Sigma-Aldrich | D9542 | 1:5000 | IF |
| Mounting medium (DABCO) | Antifade medium | Sigma-Aldrich | D2522 | N/A | IF |
| **RNA ISH reagents** |  |  |  |  |  |
| RNAscope 2.5 HD Detection Kit (Brown) | ISH detection | Advanced Cell Diagnostics | 322310 | N/A | ISH |
| *Tgfb2* probe (Mm-*Tgfb2*) | Target probe | Advanced Cell Diagnostics | 406181 | N/A | ISH |
| DapB probe | Negative control | Advanced Cell Diagnostics | 310043 | N/A | ISH |
| **Instrumentation / equipment** |  |  |  |  |  |
| Vevo 3100 + MX400 probe (30 MHz) | Echocardiography | FUJIFILM VisualSonics | Institutional | N/A | Imaging |
| Cryostat HM 525 | Frozen sectioning | Thermo Scientific | Institutional | N/A | Histology |
| Zeiss Axiovert 200 with AxioCam503 | Widefield imaging | Zeiss | Institutional | N/A | Imaging |
| Zeiss LSM 510 Meta | Confocal imaging | Zeiss | Institutional | N/A | Imaging |
| Bio-Rad CFX Connect | qPCR instrument | Bio-Rad | 1855201 | N/A | qPCR |
| NanoDrop ND-2000 | RNA quantification | Thermo Fisher Scientific | ND-2000 | N/A | RNA QC |

### Mice

All animal experiments were approved by the Institutional Animal Care and Use Committee at the University of South Carolina. Established recommendations for design, execution, and reporting studies on experimental thoracic aortic aneurysms in mice were followed^32^. Key Resources Table for detailed reagent information is provided in the Supplemental Detailed Methods. Tamoxifen-inducible, smooth muscle cell (SMC)-specific *Tgfb2* conditional knockout mice (*Tgfb2*cKO) were generated by crossing *Tgfb2^flox/flox^* mice with *Myh11CreERT2* and *ROSA*^mT/mG^ reporter strains^31, 33, 34^. Because the *Myh11CreERT*^2^ transgene is incorporated into the Y chromosome, only male mice were used. Male *Tgfb2^flox/flox^; Myh11CreER^T2+/-^*mice received intraperitoneal tamoxifen (1 mg/day for 5 days) at 4 weeks of age to induce SMC-specific deletion of *Tgfb2*. Control groups included *Tgfb2^+/+^; Myh11CreER^T2+/-^* and wild-type C57BL6J males.

### Echocardiography, Histology, qPCR, and Western Blotting Procedures

Thoracic aortic structure and function were assessed by transthoracic echocardiography (Vevo 3100®) under isoflurane anesthesia. Aortas were harvested after PBS perfusion for histology, immunofluorescence, and RNA in situ hybridization. Paraffin and frozen sections were prepared for staining with Masson’s trichrome, Verhoeff–van Gieson, and Alcian Blue, as well as for immunofluorescence using standard protocols. Aortic pathology scoring was performed as described. Gene expression was analyzed by quantitative real-time RT-PCR, and protein expression and SMAD2/3, p38 and ERK1/2 MAPK phosphorylation were evaluated by western blotting. All procedures involving physiological, pathological, and molecular characterization of the mouse model were conducted in accordance with established recommendations in the field ^32, 35^.

## Supplemental Detailed Methods

### Mice

All mouse breeding and experiments were approved by the Institutional Animal Care and Use Committee (IACUC) of the University of South Carolina (USC). Mice were bred and housed at the USC Animal Research Facility, School of Medicine. Study mice were generated by crossing male hemizygous *Myh11CreERT*^2^ transgenic mice (*Myh11CreER^T2+/-^*) heterozygous for *Tgfb2* floxed allele (*Tgfb2^+/flox^*) with female *Tgfb2^flox/flox^*mice^31, 33^. Because *Myh11CreERT*^2^ is located on the Y chromosome, only male *Myh11CreER^T2+/-^; Tgfb2f/f* mice were used. Male *Myh11CreER^T2+/-^; Tgfb2+/+* mice, wild-type C57BL6J males, and vehicle-treated *Tgfb2^+/flox^* mice served as controls. For lineage tracing, *ROSA*^mT/mG^ mice (referred to as *mTmG^+/-^* or *^-/-^*) were crossed to generate *Myh11CreERT*^2*+/-*^*; Tgfb2^flox/flox^; mTmG^+/-^* mice^34^. Male *Myh11CreER^T2+/-^; Tgfb2^+/+^; mTmG^+/-^* mice served as double fluorescent reporter controls.

*Genotyping:* Genomic DNA was extracted from embryonic tail biopsies. The *Tgfb2* floxed alleles were amplified using IMF65 (5’-CACCTTTTACCTACAGATGAAGTTGC-3’), IMR65 (5’-CTTAAGACCACACTGTGAGATAATCC-3’), and IMR66 (5’-CAACGGGTTCTTCTGTTAGTCC-3’) primer pairs, respectively, as previously described^31^.

*Myh11-CreER^T2^*: IMR7338 – 5’-CTAGGCCACAGAATTGAAAGATCT-3’, IMR7339- 5’- GTAGGTGGAAATTCTAGCATCATCC-3’, IMF104 - 5’-TGA CCC CAT CTC TTC ACT CC- 3’, IMR105 – 5’- 5’AGT CCC TCA CAT CCT CAG GTT-3’

*mT/mG^+/-^*reporter allele: WTF - 5’-CTC TGC TGC CTC CTG GCT TCT-3’, WTR - 5’-CGA GGC GGA TCA CAA GCAATA-3’ and MuR - 5’-TCA ATG GGC GGG GGT CGT T-3’.

*Tamoxifen treatment:* At 4 weeks of age, mice were injected intraperitoneally with tamoxifen (TX) (T5648, Sigma-Aldrich) (1 mg/d i.p.) for 5 consecutive days to induce SMC-specific *Tgfb2* deletion. Tamoxifen solution was prepared by dissolving 40 mg tamoxifen in 400 µL ethanol (heated and vortexed), followed by addition of 3.6 mL corn oil (Sigma C8267), vortexing, and probe sonication. *Tgfb2* wild-type controls were injected with corn oil vehicle (C8267, Sigma-Aldrich).

### Echocardiography

Transthoracic echocardiography was performed using a Vevo 3100® system with MX400 probe (30 MHz; FUJIFILM VisualSonics, Toronto, Canada) as described ^32, 35^. Mice were anesthetized with 3% isoflurane for induction and maintained at 1.5% via nose cone. Body temperature was stabilized on a heated platform, and eyes protected with ophthalmic ointment. Chest and abdomen hair were removed with Nair™ lotion, wiped with water-soaked gauze, and electrode gel was applied to leads for ECG monitoring. Heart rate was maintained between 450–550 bpm. Ultrasound gel was applied, and parasternal long-axis images were acquired in B-, M-, and Doppler-modes. Data was analyzed using NIH ImageJ and Vevo Lab software.

### Tissue Collection and Processing

Tissue collection and processing was done in accordance with established recommendations in the field ^32^. Briefly, aortas from *Tgfb2*cKO and control mice were harvested following PBS perfusion through the left ventricle. Tissues were fixed in 4% paraformaldehyde (PFA) overnight at 4°C for histology. For paraffin embedding, samples were dehydrated in graded alcohols, cleared in xylene, and embedded in paraffin; 7 µm sections were prepared for histology, immunohistochemistry, and RNAscope in situ hybridization. For frozen sections, PFA-fixed aortas were cryoprotected in 30% sucrose overnight, rinsed in PBS, infiltrated with OCT, oriented, snap-frozen in chilled isopentane, and stored at –80°C. Ten-µm cryosections were cut using a Thermo Scientific MICROM HM 525 cryostat. For western blotting and zymography, aortas were snap-frozen in liquid nitrogen after removal of perivascular fat and stored at –80°C.

### Histology

Histological staining and examination was done in accordance with established recommendations in the field ^32^.Paraffin sections were deparaffinized in xylene, rehydrated in graded alcohols, and stained with Von Kossa/Alcian Blue, Masson’s trichrome, and Verhoeff– van Gieson stains (American MasterTech, StatLab, TX) to evaluate extracellular matrix remodeling. Aortic pathology was scored as described^36^.

*Alcian Blue Staining:* After deparaffinization, sections were immersed in 3% acetic acid for 3 min, stained with Alcian Blue (pH 2.5) for 30 min, rinsed, counterstained with Nuclear Fast Red for 5 min, dehydrated, cleared in xylene, and mounted (Vector Lab H5000).

*Masson’s Trichrome:* Sections were deparaffinized, hydrated, mordanted in Bouin’s fluid (1 h, 56°C), rinsed, stained with Weigert’s hematoxylin (5 min), Biebrich Scarlet–Acid Fuchsin (15 min), phosphomolybdic/phosphotungstic acid (15 min), and Aniline Blue (10 min). Slides were differentiated in 1% acetic acid (3 min), dehydrated, cleared in xylene, and mounted.

*Verhoeff–van Gieson Elastic Stain:* Sections were stained with Verhoeff’s elastic solution (15 min), differentiated in 2% ferric chloride (3 min), treated with 5% sodium thiosulfate (1 min), counterstained with van Gieson’s solution (30 s), dehydrated, cleared in xylene, and mounted.

### Immunofluorescence

Both paraffin and frozen sections were used. For paraffin sections, antigen retrieval was performed by heating in citrate buffer (pH 6, Vector H3300-250). For frozen sections, antigen retrieval was performed with proteinase K (1:100, Millipore JA1477, 20 min). After rinsing in PBS, slides were blocked (5% normal serum + 1% BSA + 0.1% Tween-20 in PBS, 20 min), incubated overnight with primary antibodies at 4°C, washed, and incubated with fluorescent secondary antibodies for 30 min. Nuclei were counterstained with DAPI (5 min), and slides mounted in DABCO (Sigma). Double reporter tissues were mounted in glycerol:methanol (1:1) to reduce endogenous GFP and tdTomato signals.

### Cell Fate Mapping

For lineage tracing, *ROSA*^mT/mG^ reporter mice (called as *mTmG*^+/-^ or *mTmG*^-/-^) were used, in which Cre-mediated recombination induces a switch from membrane-targeted Tomato (mT, red) to membrane-targeted GFP (mG, green) expression^34, 37^. These mice were crossed with *Myh11CreER*^T2+/-^; *Tgfb2*^flox/flox^ mice to generate experimental animals with the genotype *Myh11CreER*^T2+/-^; *Tgfb2*^flox/flox^; *mTmG*^+/-^. To induce recombination in smooth muscle cells (SMCs), tamoxifen was administered. *Myh11CreER*^T2+/-^; *Tgfb2*^+/+^; *mTmG*^+/-^ mice served as controls to evaluate baseline reporter expression and recombination efficiency in the absence of *Tgfb2* deletion. Frozen sections were air-dried, washed in PBS, stained with DAPI, mounted in DABCO, and imaged using Zeiss Axiovert 200 with AxioCam503 and Zeiss LSM 510 Meta confocal microscope (Oberkochen, Germany).

### RNA in situ hybridization

Detection of *Tgfb2* transcripts was performed using the RNAscope® 2.5 HD Detection Kit (Brown; Cat# 322310, Advanced Cell Diagnostics, Newark, CA). Embryos (E12.5–13.5) were hybridized with a *Tgfb2*-specific probe (Mm-*Tgfb2*; Cat# 406181, ACD), and a bacterial DapB probe (Cat# 310043, ACD) served as a negative control. For pretreatment, slides underwent heat-mediated antigen retrieval in a microwave for 15 min, followed by protease digestion for 30 min at 40 °C in a HybEZ™ oven. After hybridization and amplification, sections were counterstained with hematoxylin (1:10 dilution, 1 min). Images were collected at 40× magnification from 4–5 randomly selected regions per section.

Quantitative analysis was performed in Fiji (NIH) using the workflow described in ACD Technical Note TS 46-003. Nuclei (≥500 pixels², circularity 0.25–1) were automatically segmented, and at least 500 nuclei per animal were evaluated. *Tgfb2* probe signals were identified with the “Weka Segmentation” plugin (Fast Random Forest classifier) using thresholds of ≥1 pixel² and circularity 0.25–1. Probe clusters were counted as single events regardless of dot density. At least three embryos were analyzed for each experimental group.

### Quantitative PCR

RNA isolation was done using Qiagen miRNeasy® Mini kit according to the manufacturer’s instruction (QIAGEN GmbH, QiAGEN, Hilden, Germany).^38^ Briefly, 10-15 mg thoracic aortic tissue without perivascular adipose tissue was placed in 2 mL centrifuge tubes and 700 mL QIAzol® lysis reagent and a 5 mm stainless steel bead were added to each tube. The tissue was homogenized in Qiagen TissueLyser II. After the homogenate was incubated at RT for 5 minutes, 140 µL chloroform solution was added and it was shaken vigorously for 15 seconds. Then the homogenate was incubated at RT for 3 minutes and centrifuged for 15 min at 12,000X g at 4°C. Next, the upper aqueous phase was transferred to a new 1.5 mL collection tube and 525 mL 100% ethanol was added mixed thoroughly by pipetting. A sample aliquot (700 µL) was pipetted into an RNeasy® Mini column in a 2 mL collection tube and centrifuged for 15 seconds (s) at 12,000X g at RT. After discarding the flowthrough, DNase digestion was done by adding 80 µL DNase I incubation mix directly to the RNeasy spin column membrane and placed it for 15 min at RT. Then, 700 µL of Buffer RWT was added to the RNeasy Mini column and centrifuged for 15 s at 12,000X g at RT. Next, 500 µL Buffer RPE was added to the RNeasy spin column and centrifuged for 15 s at 12,000X g at RT. After discarding the flowthrough, another 500 µL Buffer RPE was added to the RNeasy spin column and centrifuged for 2 min at 12,000X g at RT. The RNeasy spin column was then transferred to a new 1.5 mL collection tube, 30 µL RNase free water was pipetted directly onto the column membrane and centrifuged for 2 min at 12,000X g at RT to elute.

Quantification of RNA was done using NanoDrop Microvolume Spectrophotometer (ThermoFisher Scientific) and complementary DNA was synthesized from 300 ng RNA using BIO-RAD iScript™ Reverse Transcription Supermix for RT-qPCR (cat # 1708841). The PCR product was diluted with RNase free water (1:5) for qPCR analysis. To perform qPCR 10 µL iTaq Universal SYBR® Green supermix (BioRad Cat # 1725124), 1 µL primer, 4 µL RNase free water and 5 µL cDNA were added into wells of PCR plate (BioRad Cat #HSP9601). The plate was sealed with a plate sealer and run on BioRad CFX connect system using the following protocol. Initial melting at 94 °C for 3 minutes and 40 cycles of 95 °C for 15 sec, 60 °C for 20 sec and 72 °C for 15 seconds. Real-time qPCR amplification was performed using BioRad Prime PCR assay. The following primers: smooth muscle myosin heavy chain (*Myh11*) (qMmuCID0019272), SM α-actin (*Acta2*) (qMmu CID0006375), *Tgfb1* (qMmuCED0044726), *Tgfb2* (qMmuCID0024408), *Tgfb3* (qMmuCID0045203), *Tgfbr1*(qMmuCID0026554), *Tgfbr2* (qMmuCID0015359), Elastin (*Eln*) (qMmuCID0006420), lysyl oxidase (*Lox*) (qMmuCID0023621), and *Gapdh* (qMmuCED0027497) from were used (BIO-RAD Hercules, CA).

### Western Blotting

Western blotting was done in accordance with established procedures and recommendations in the field ^32, 39^. Protein samples were stored at –80 °C until analysis. Equal amounts of protein were separated by SDS–PAGE and transferred to PVDF membranes.

Membranes were probed with primary antibodies (all from Cell Signaling Technology, Danvers, MA) against phospho-SMAD2 (#3108), SMAD2 (#5339), phospho-SMAD3 (#9520), SMAD3 (#9523), phospho-SMAD1/5 (#9516), SMAD1/5 (#9743), phospho-p38 (#4511), p38 (#8690), phospho-ERK1/2 (#4370), and ERK1/2 (#4695), each at 1:1000 dilution. β-actin (clone AC-15, Sigma-Aldrich, St. Louis, MO) was used as a loading control. Horseradish peroxidase– conjugated secondary antibodies (anti-mouse or anti-rabbit IgG, #7074, Cell Signaling Technology) were applied at 1:5000 dilution. Bands were visualized using Clarity™ ECL reagents (Bio-Rad Laboratories, Hercules, CA) and captured on X-OMAT AR film (Eastman Kodak, Rochester, NY). Films were scanned (Epson) and images processed with Adobe Photoshop (Adobe Systems, Seattle, WA).

Densitometric analysis was performed using NIH ImageJ (Fiji). Phosphorylated protein levels were normalized to their respective total protein or to β-actin. Data were recorded in Microsoft Excel and analyzed using GraphPad Prism 9 (GraphPad Software, San Diego, CA). Results are expressed as mean ± SD. Comparisons between groups were performed using one-way ANOVA or unpaired two-tailed Student’s t test, with P < 0.05 considered statistically significant.

## Results

### Generation of Mice with Postnatal Smooth Muscle Cell-Specific *Tgfb2* Deletion in the Thoracic Aorta

Because *Tgfb2* knockout fetuses die perinatally, we induced *Tgfb2* deletion postnatally at 4 weeks of age in SMCs (Figure 1A-B). Homozygous *Tgfb2^flox/flox^* mice, in which exon 2 is flanked by loxP sites, were crossed with mice expressing tamoxifen (TX)-inducible Cre-ER^T2^ under the control of the SMC-specific *Myh11* promoter (*Myh11CreER*^T2^). This cross generated experimental *Myh11CreER^T2^; Tgfb2^flox/flox^* mice (hereafter referred to as *Tgfb2*cKO) **(Figure 1 B).** To induce recombination, animals were injected intraperitoneally with 1 mg of tamoxifen daily for five consecutive days. TX-treated *Myh11CreER^T2^; Tgfb2^+/+^* mice and vehicle-injected *Myh11CreER^T2^; Tgfb2^flox/flox^* mice served as controls, as their aortic histology was normal and indistinguishable. Conditional deletion of *Tgfb2* in SMCs was first confirmed by PCR genotyping of thoracic aortas (with perivascular adipose tissue removed) 4-weeks after TX treatment, demonstrating specific identification of the deleted *Tgfb2* allele in *Tgfb2*cKO mice^31^. Control mice showed no *Tgfb2* deletion. To further validate the efficiency and specificity of TX-induced Cre recombination, *Myh11CreER^T2^; Tgfb2^flox/flox^* mice were intercrossed with *mT/mG*^+/-^ double-fluorescent reporter mice. Both heterozygous (*mTmG^+/-^*) and homozygous (*mTmG^-/-^*) reporters were used. Following TX injection, Cre recombinase became activated in SMCs, entered the nucleus, and excised both the tdTomato (mT) cassette and exon 2 of *Tgfb2*.

**Figure 1.**
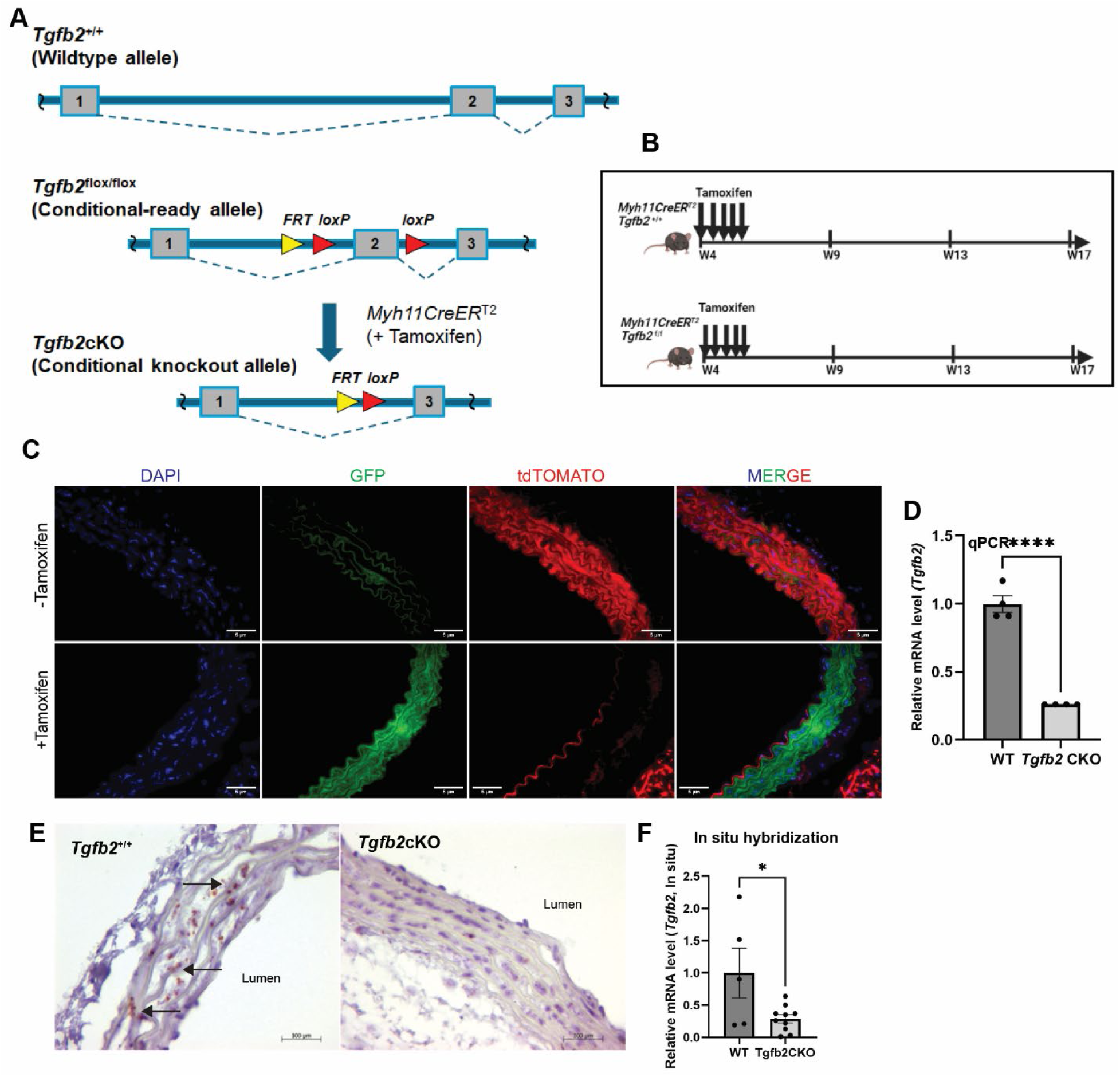
Generation and validation of SMC-specific *Tgfb2* conditional knockout mice. (A) Schematic of Cre-mediated removal of exon 2 of *Tgfb2*. (B) Experimental design: *Myh11CreER*^T2^;*Tgfb2*^flox/flox^ mice were injected intraperitoneally with tamoxifen at 4 weeks of age to delete *Tgfb2* in vascular smooth muscle cells (SMCs). (C) Double-fluorescent reporter mice (*Myh11CreER*^T2^;*Tgfb2*^flox/flox^;*mT/mG^+/-^*) showing recombination efficiency and specificity at 4 weeks post-TX (9 weeks of age): SMCs express membrane GFP, while non-SMCs express membrane tdTomato (RFP). (D) Quantitative real-time PCR of aortic tissue lysates from 9-weeks-old mice demonstrating reduced *Tgfb2* expression in conditional knockout (cKO) mice. (C) RNAscope in situ hybridization of *Tgfb2* in the aortic media of 9-weeks-old wild-type (black arrows) and *Tgfb2* cKO mice, showing marked reduction in *Tgfb2* expression. (F) Quantification of *Tgfb2* RNAscope signal. Scale bar: C, 5 µm; E, 20 µm. Data shown as mean ± SEM. *P < 0.05; ***P < 0.001. Abbreviations: Flox, floxed allele.

Expression of membrane EGFP (mG) exclusively in SMCs of the aortic media confirmed effective Cre-mediated *Tgfb2* deletion in *Myh11CreER^T2^; Tgfb2^flox/flox^; mTmG^+/-or-/-^* mice. Control animals expressed only tdTomato (mT) in their ascending and descending thoracic aortas **(Figure 1C)**. Quantitative RT-PCR (qPCR) demonstrated significant downregulation of *Tgfb2* expression in thoracic aortas of *Tgfb2*cKO mice relative to controls **(Figure 1D)**. These results were validated and extended by RNAscope in situ hybridization, which revealed significant *Tgfb2* expression in medial SMCs of control mice, but significantly reduced *Tgfb2* signal in *Tgfb2*cKO mice **(Figure 1E-F)**.

### Postnatal SMC-Specific *Tgfb2* Deletion Causes Thoracic Aortic Aneurysm

The incidence of thoracic aortic aneurysm was evaluated in vivo at 9 weeks of age (i.e., 4-weeks after TX-induced *Tgfb2* deletion) using transthoracic echocardiography (Vevo 3100 with MX400 probe, 30 MHz) in parasternal long-axis view. Serial imaging demonstrated progressive dilation of both ascending and descending aortas in *Tgfb2*cKO mice compared with controls **(Figure 2A– F)**. The ascending aortic diameter was significantly greater in *Tgfb2*cKO mice than in controls (1.695 ± 0.0525 mm vs 1.475 ± 0.038 mm, *P* = 0.0016) **(Figure 2A–C)**. The descending aorta was similarly enlarged (1.265 ± 0.0419 mm vs 1.05 ± 0.0423 mm, *P* = 0.0009) **(Figure 2D–F)**.

**Figure 2.**
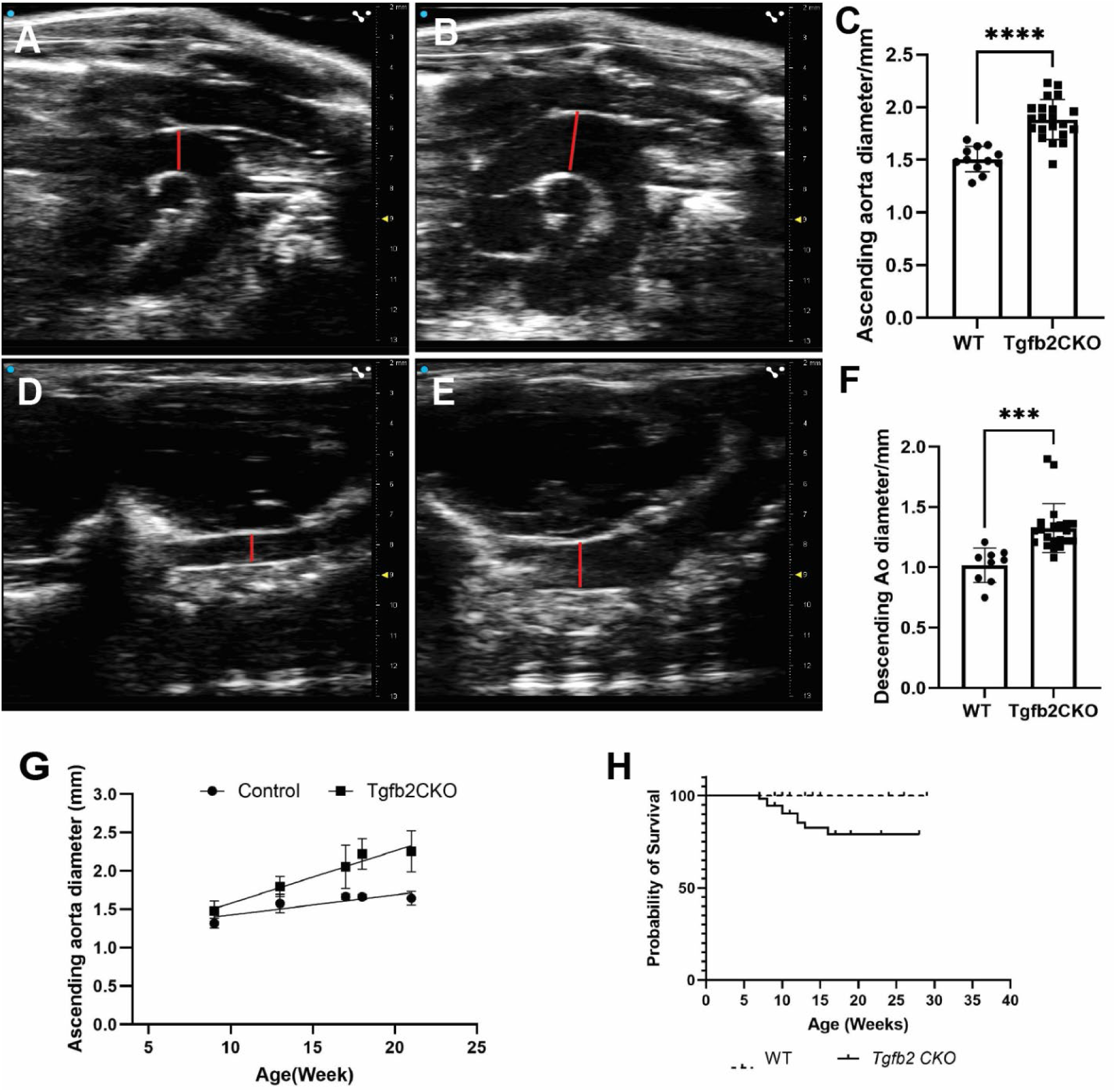
SMC-specific deletion of *Tgfb2* leads to dilation of the ascending and descending aorta. Transthoracic echocardiography was performed beginning 4 weeks after completion of tamoxifen injections (9 weeks of age) using a Vevo 3100® system with an MX400 probe (30 MHz). Representative images of the ascending aorta from wild-type (WT) mice **(A)** and SMC- *Tgfb2* inducible conditional knockout (cKO) mice **(B)**, quantified in **(C)**. Representative images of the descending aorta from WT mice **(D)** and *Tgfb2*cKO mice **(E)**, quantified in **(F)**. **(G)** Time- course analysis showed a significant increase in ascending aortic diameter in *Tgfb2*cKO mice compared with WT (P = 0.0081). **(H)** Kaplan–Meier survival curve demonstrated ∼15% mortality from aortic rupture in *Tgfb2*cKO mice, in some cases as early as 2 weeks post-deletion. Internal diameters were measured at maximal dilation from echocardiogram video clips using NIH ImageJ or VevoLab software. Data are mean ± SEM. ***P < 0.001; ****P < 0.0001.

By 13 weeks of age (i.e., 8-weeks post-TX), ascending aortas of *Tgfb2*cKO mice reached an average of 1.883 ± 0.0418 mm compared to 1.507 ± 0.035 mm in controls (*P* = 0.0009), while descending aortas measured 1.325 ± 0.0442 mm versus 1.017 ± 0.0475 mm (*P* = 0.0003).

Maximal dilation was observed in the proximal descending aorta, occasionally reaching 2.0 mm **(Figure 2F)**. Longitudinal follow-up of the TAA confirmed age-dependent progression aneurysm in *Tgfb2*cKO mice **(Figure 2G)**. After echocardiography, mice were examined grossly for dissection or hemorrhage. In a survival study of 60 *Tgfb2*cKO and 60 control animals followed to 30 weeks, 10 *Tgfb2*cKO mice died, while no deaths occurred in controls **(Figure 2H, Survival Curve)**.

### Postnatal Loss of SMC-Specific *Tgfb2* Causes Thoracic Aortic Dissection and Rupture

Histological analysis of serial hematoxylin and eosin–stained sections revealed that *Tgfb2* deletion induced aneurysm and dissection as early as 7 weeks of age (i.e., two weeks after TX treatment). Among 60 *Tgfb2*cKO mice, 10 (16.7%) died of internal hemorrhage from aortic rupture. Postmortem analysis showed rupture at the proximal descending aorta in 8 mice and at the distal descending aorta in 2 mice **(Figure 3A-B)**. Neither TX-injected *Myh11CreER^T2^; Tgfb2^+/+^* mice nor vehicle-injected *Myh11CreER^T2^; Tgfb2^flox/flox^* mice showed rupture or hemorrhage. Ruptures occurred between 7 and 15 weeks of age, with 80% of cases dying within 8 weeks after deletion. Gross and microscopic examination showed intramural hematoma and ulceration in the ascending aorta and arch, and aortic dissections in the descending aorta, most prominently in the proximal descending segment **(Figure 3 C-D)**.

**Figure 3.**
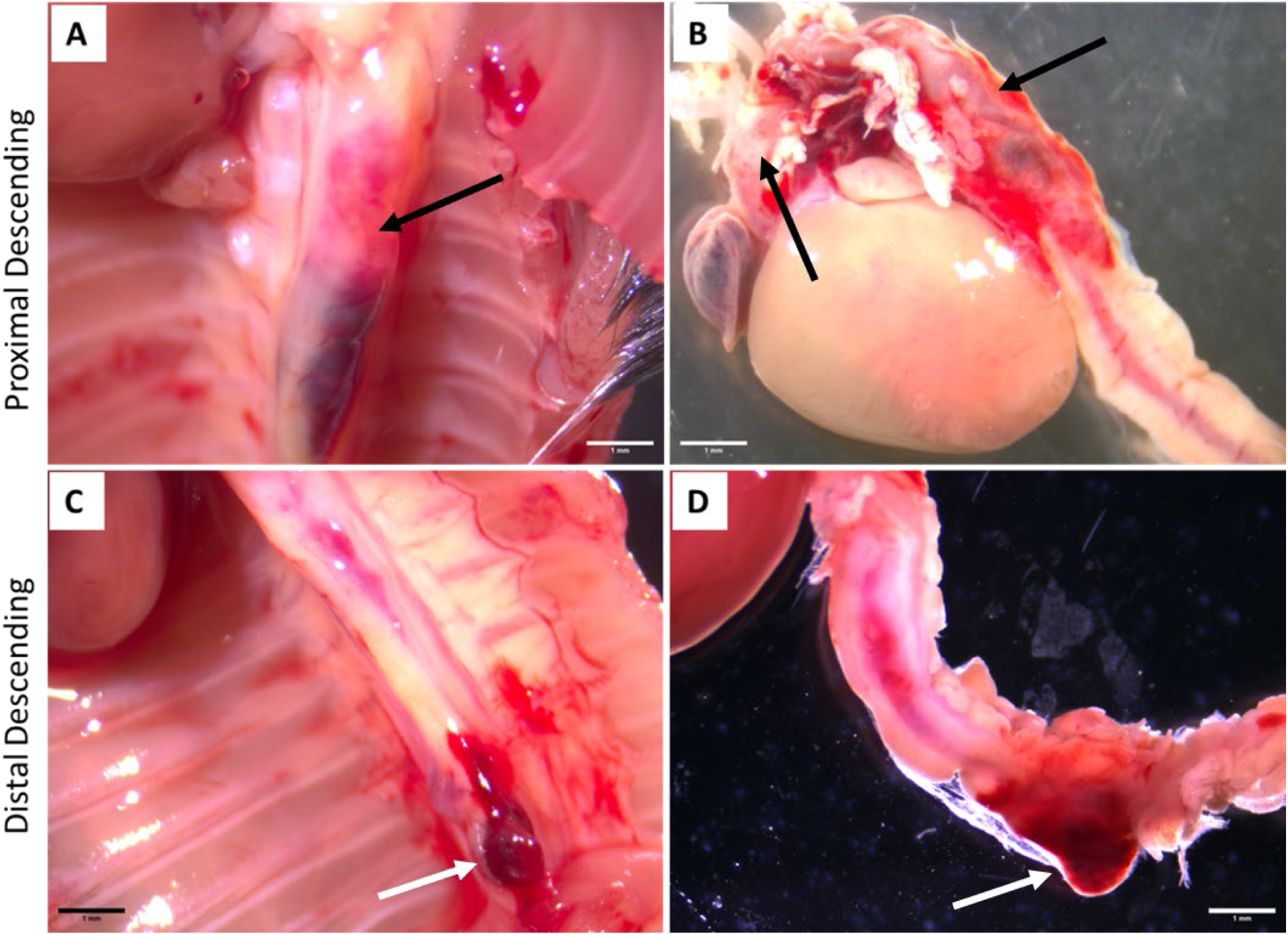
Loss of *Tgfb2* in SMCs results in intramural hematoma, dissection, and aortic rupture. Gross examination of aortic pathology in mice at 7 weeks of age (i.e., two weeks after TX treatment). SMC-*Tgfb2* inducible conditional knockout mice developed intramural hematomas (arrows). **(A, B)** Representative images of hematomas in the ascending aorta, aortic arch, and proximal descending aorta (black arrows). **(C, D)** Representative images of intramural hematoma and dissection in the distal descending aorta (white arrows). Scale bar = 1 mm.

### SMC-Specific *Tgfb2* Loss Leads to Medial Degeneration, Adventitial Thickening, and Excess ECM Deposition

Histopathological assessment at 9 weeks of age (i.e., 4-weeks after TX induction) demonstrated marked abnormalities across all three layers of the *Tgfb2*cKO aortic wall. Both ascending and descending thoracic aortas displayed medial degeneration, medial thickening, adventitial enlargement, and occasional intimal denudation **(Figure 4 A–F)**. The severity of pathology varied by location: elastic fiber fragmentation, dissection, and adventitial thickening were more prominent in the descending aorta than in the ascending segment, especially in younger mice **(Figure 4 G–J)**. Medial degeneration ranged from mild elastic fiber fragmentation to near-complete breakdown of elastic lamellae, with SMCs losing their characteristic elongated nuclear morphology and circumferential alignment and appeared significantly stretched **(Figure 4 K, L-M, N)**. Approximately 32% (19/60) of animals developed severe medial degeneration within four weeks of gene deletion. In mice without dissection, moderate but progressive elastic fragmentation was observed, accompanied by marked lamellar widening and highly stretched, irregular, and abnormal shape of SMC **(Figure 4 L–M, N)**. Nearly 42% (25/60) of *Tgfb2*cKO mice developed false lumens within the descending aorta. These often originated in the proximal descending portion and re-entered the true lumen at varying distal points, with some extending toward the distal thoracic aorta. Penetrating ulcers exposing the inner media were frequently identified in the ascending aorta, whereas in 6/60 mice the intimal layer and internal elastic lamina detached and folded into the lumen leading to an intimal flap in the descending aorta.

**Figure 4.**
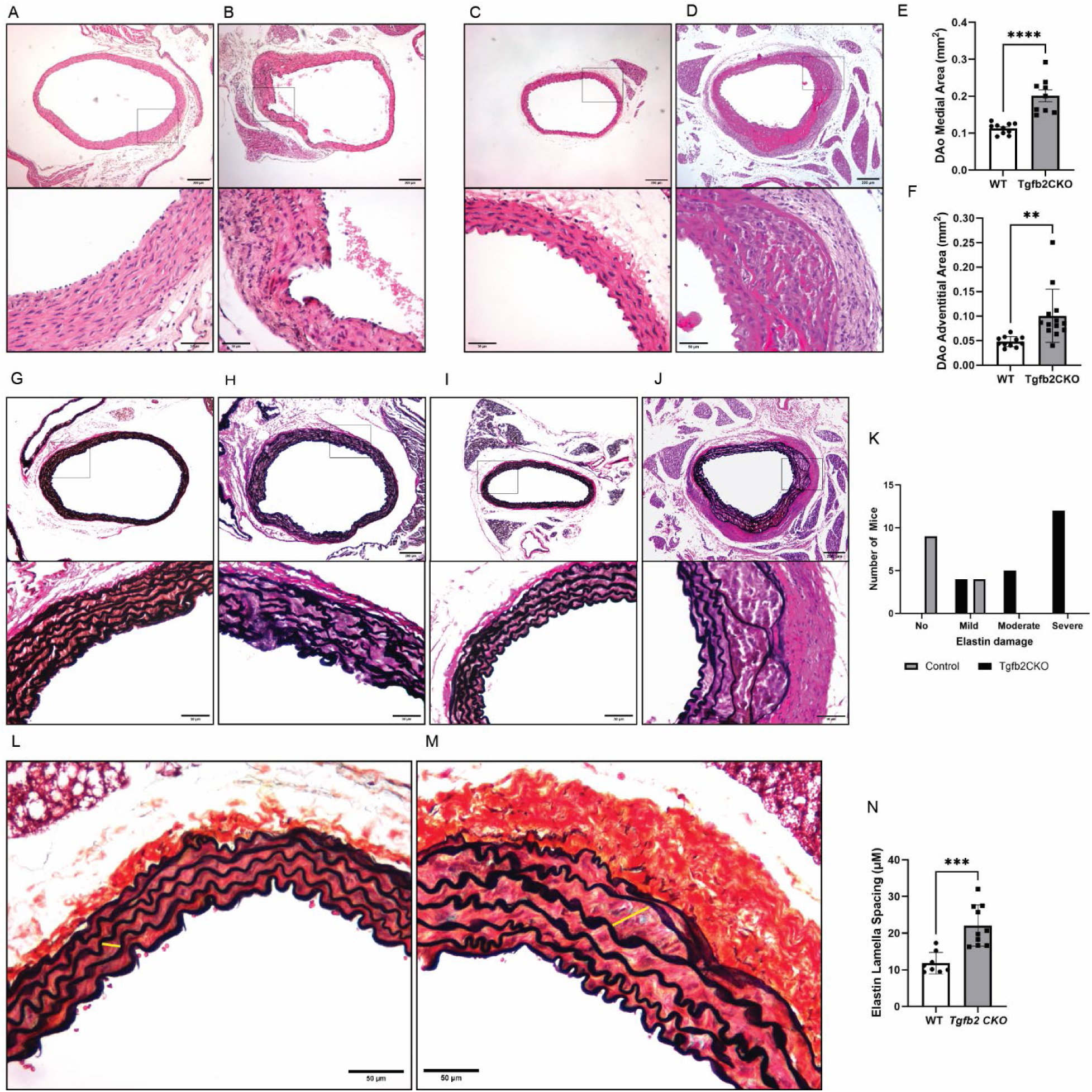
Loss of SMC-TGFβ2 is associated with histological changes in the aortic wall. Histological analyses at 9 weeks of age (i.e., 4-weeks after TX treatment). **(A)** Representative H&E images of the ascending aorta from wild-type (WT) mice with magnified boxed region. **(B)** Ascending aorta from *Tgfb2* conditional knockout (cKO) mice showing an intimal tear (magnified boxed region). **(C)** Descending aorta from WT mice with magnified boxed region. **(D)** Descending aorta from *Tgfb2*cKO mice showing aortic dissection, medial widening, and adventitial thickening, quantified in **(E, F). (G, H)** Verhoeff–Van Gieson (VVG) elastin staining of the ascending aorta from WT **(G)** and *Tgfb2*cKO **(H)** mice with magnified regions; *Tgfb2*cKO mice show elastin fiber degradation. **(I, J)** VVG staining of the descending aorta from WT **(I)** and *Tgfb2*cKO **(J)** mice; *Tgfb2*cKO mice show elastin breaks and dissection. **(K)** Chi-square analysis of elastin damage in WT vs *Tgfb2*cKO groups. **(L, M)** Pentachrome staining of descending aorta from WT **(L)** and *Tgfb2*cKO **(M)** mice, showing increased elastin lamellar spacing (yellow lines), quantified in **(N)**. Data are mean ± SEM. **P < 0.01; ***P < 0.001; ****P < 0.0001.

Histological sections revealed the presence of red blood cells and thrombi as intramural hematoma within the dissected medial wall and false lumen spaces **(Figure 4 D, J)**.

Masson’s Trichrome staining demonstrated extensive collagen accumulation in both medial and adventitial layers of *Tgfb2*cKO aortas, affecting both ascending and descending regions **(Figure 5 A–E)**. Collagen was particularly concentrated in zones of medial degeneration and dissection, while adventitial deposition contributed substantially to wall thickening. Alcian Blue staining further revealed abnormal proteoglycans accumulation in dissected medial regions and false lumens compared with controls **(Figure 5 F–H)**. Finally, immunohistochemical analysis using Ki-67 as a proliferation marker showed increased aortic cell proliferation in the medial wall of *Tgfb2*cKO mice **(Figure 5 I–K)**.

**Figure 5.**
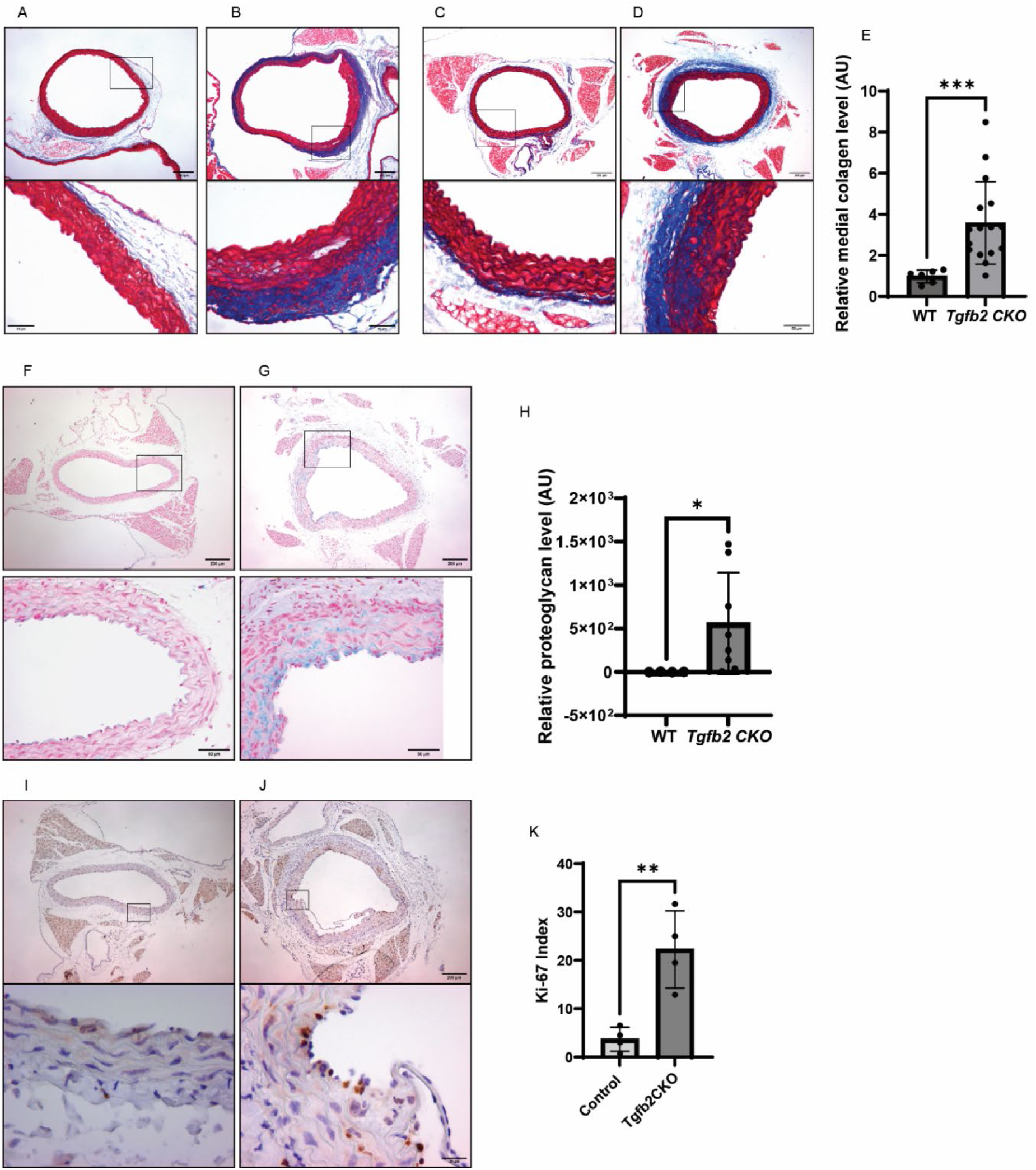
SMC-*Tgfb2* deletion results in collagen and proteoglycan deposition and increased cell proliferation in the aortic wall. Histological analyses at 9 weeks of age (i.e., 4- weeks after TX treatment). **(A, B)** Masson’s Trichrome staining of the ascending aorta from wild-type (WT) **(A)** and *Tgfb2* conditional knockout (cKO) mice **(B)**, showing increased collagen deposition (blue) in cKO mice. **(C, D)** Masson’s Trichrome staining of the descending aorta from WT **(C)** and *Tgfb2*cKO mice **(D)**, with magnified regions showing elevated collagen deposition in the media and adventitia. **(E)** Quantification of collagen deposition by unpaired two-tailed t test. **(F, G)** Alcian blue staining of descending aorta from WT **(F)** and *Tgfb2*cKO **(G)** mice showing increased proteoglycan deposition in the medial layer. **(H)** Quantification of proteoglycan deposition. Collagen and proteoglycans were quantified in NIH ImageJ using the color deconvolution plugin for Masson’s Trichrome and Alcian blue, respectively; mean pixel values (gray scale) of blue staining after thresholding were used for histograms. **(I, J)** Immunohistochemistry for Ki67 in the descending aorta from WT **(I)** and *Tgfb2*cKO **(J)** mice, showing increased proliferating cells (brown nuclei) in cKO mice. **(K)** Quantification of Ki67- positive nuclei. Data are mean ± SEM. **P = 0.01; ***P < 0.001.

### Loss of TGFβ2 in SMCs Alters Gene Expression Linked to Differentiation and ECM Remodeling

Quantitative PCR analysis of thoracic aortas (including ascending, arch, and descending regions with perivascular fat removed) demonstrated pronounced transcriptional alterations in *Tgfb2* inducible conditional knockout mice at 2 weeks after induction of *Tgfb2* deletion (7-weeks-old mice). Expression levels of *Tgfb2* and *Tgfb3* were significantly decreased, while *Tgfb1* and *Tgfbr2* were markedly upregulated compared with controls **(Figure 6 A–E)**. Genes associated with extracellular matrix (ECM) organization, including *Eln* (tropoelastin) and *Lox* (lysyl oxidase), were also significantly downregulated **(Figure 7 F–G)**. Furthermore, smooth muscle contractile genes *Acta2* (α-smooth muscle actin) and *Myh11* (smooth muscle myosin heavy chain) showed marked reduction in *Tgfb2*cKO mice relative to controls **(Figure 7 H–I)**.

**Figure 6.**
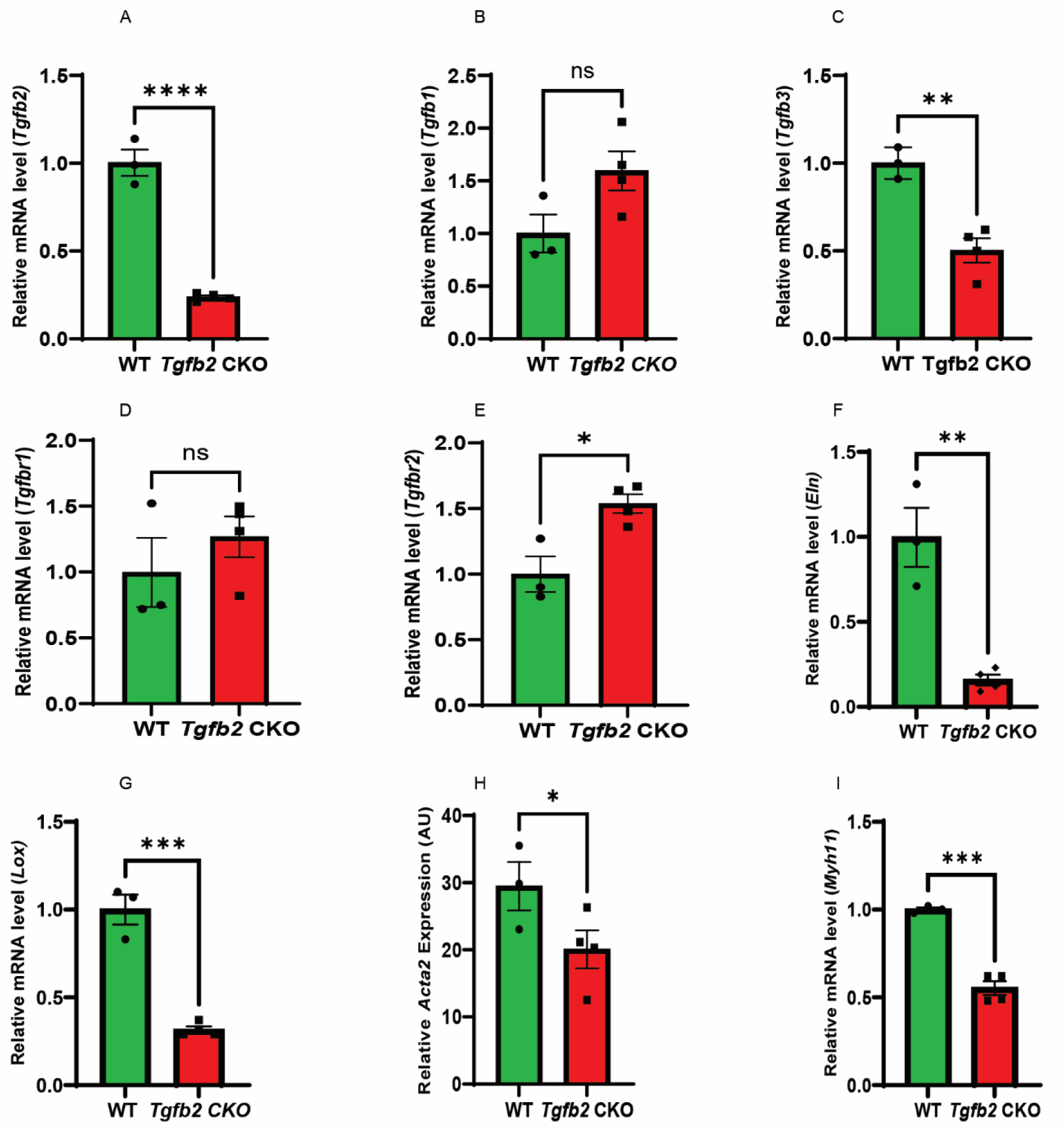
SMC-specific loss of TGFβ2 alters expression of SMC- and ECM-related genes. Real-time qPCR analysis of aortic tissue from wild-type (WT) and *Tgfb2* inducible conditional knockout (cKO) mice 2 weeks after induction of *Tgfb2* deletion (7-weeks-old). **(A)** Reduced *Tgfb2* expression. **(B)** Modest increase in *Tgfb1* expression (P = 0.0763). **(C)** Significant reduction in *Tgfb3* expression. **(D)** No difference in *Tgfbr1* expression. **(E)** Increased *Tgfbr2* expression. **(F)** Reduced elastin expression. **(G)** Reduced *Lox* expression. **(H)** Significant reduction in *Acta2* expression (P<0.05). **(I)** Significant reduction in *Myh11* expression. Gene expression was normalized to GAPDH and expressed relative to WT controls. Data are mean ± SEM. *P < 0.05; **P < 0.01; ***P < 0.001.

**Figure 7.**
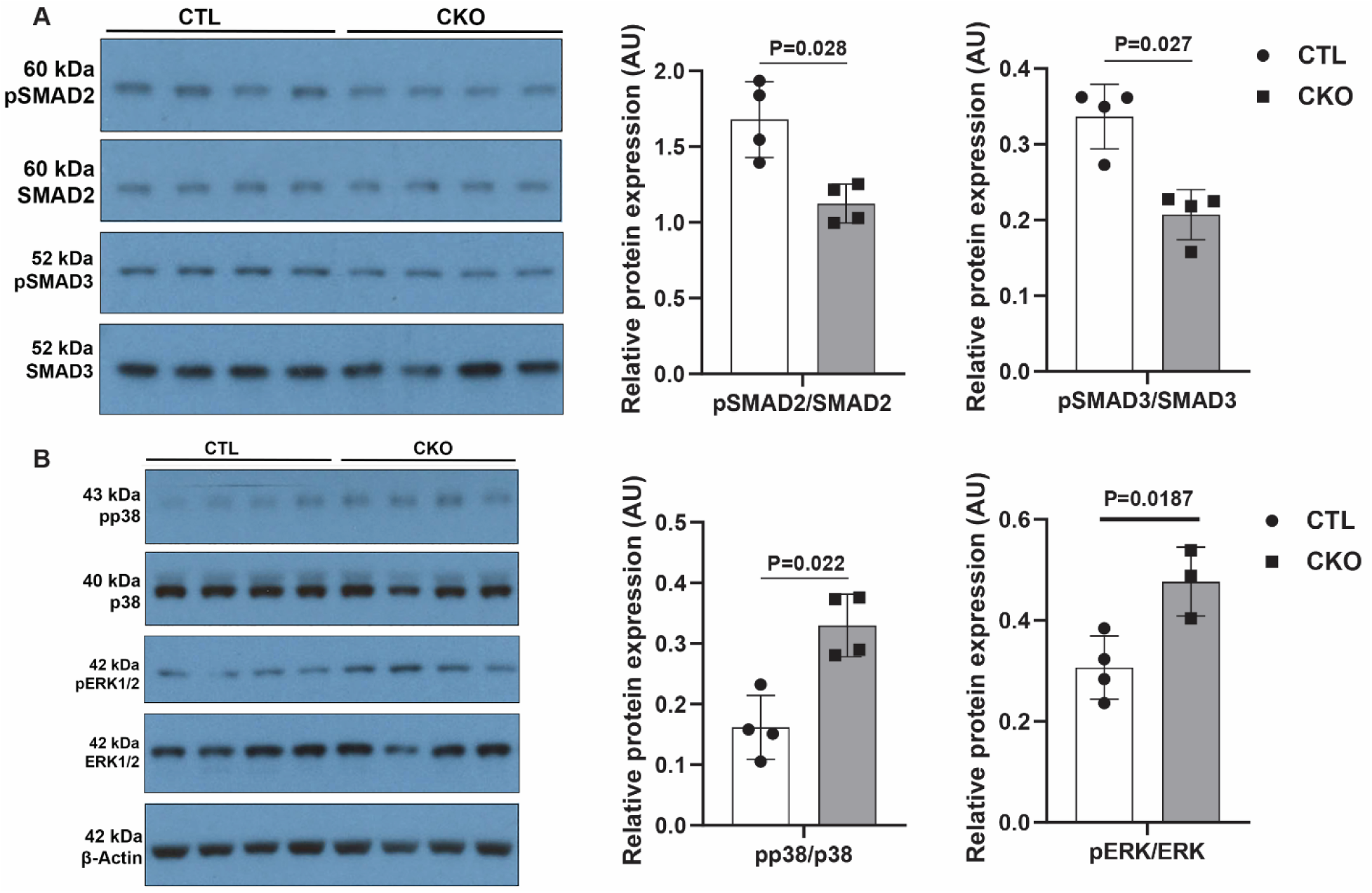
*Tgfb2* is required for maintaining physiological TGFβ signaling in the postnatal thoracic aorta. Western blot analysis of aortic tissue 2 weeks after tamoxifen induction (7 weeks of age) showing reduced phosphorylation of SMAD2 and SMAD3, and increased phosphorylation of p38 and ERK1/2 MAP kinases in SMC-*Tgfb2* inducible conditional knockout (cKO) mice compared with wild-type (WT) controls. Histograms of pSMAD2/total SMAD2, pSMAD3/total SMAD3, pp38/total p38, and pERK1/2/total ERK1/2 confirm the western blot findings.

Collectively, these transcriptional changes suggest that loss of TGFβ2 disrupts both SMC contractile identity and extracellular matrix stability, thereby predisposing the aortic wall to degeneration and aneurysm formation.

### SMC-specific *Tgfb2* deletion disrupts TGFβ–SMAD signaling

To assess early effects of *Tgfb2* loss on canonical TGFβ signaling, western blotting was performed on thoracic aortas (without perivascular adipose tissue) at 7 weeks of age, 2 weeks after tamoxifen induction. Phosphorylated SMAD2 (pSMAD2) and SMAD3 (pSMAD3) levels were significantly reduced in *Tgfb2*cKO mice, whereas total SMAD2/3 levels remained unchanged compared with controls **(Figure 7A)**. In contrast, *Tgfb2* deletion in SMCs was associated with a marked increase in phosphorylation of p38 and ERK1/2 MAPKs **(Figure 7B)**. These findings demonstrate that loss of *Tgfb2* suppresses canonical TGFβ–SMAD signaling while simultaneously enhancing non-canonical MAPK signaling during the early stages of aneurysm development.

## Discussion

In this study, we employed a tamoxifen-inducible, smooth muscle cell (SMC)-specific *Myh11CreER*^T2^ driver to delete *Tgfb2* in mice carrying a conditional allele at 4 weeks of age. Postnatal *Tgfb2* deletion in vascular SMCs produced a severe aortopathy, with aneurysm formation, dissection, and frequent sudden death from rupture. These mice developed progressive, time-dependent dilation of both ascending and descending aortas. Intramural hematomas were observed throughout the ascending aorta, aortic arch, and descending thoracic aorta, with the proximal descending region most consistently affected. Histological evaluation revealed extensive medial degeneration, medial thickening, and adventitial remodeling, accompanied by abundant collagen and proteoglycan deposition.

Our findings extend prior genetic studies of *Tgfb2* deficiency. Complete loss of *Tgfb2* results in perinatal lethality with congenital cyanosis and malformations of the heart and great vessels, including a dilated, hypercellular ascending aorta at E18.5^31^. Haploinsufficiency, by contrast, allows survival but predisposes to dilation of the aortic annulus and root with preservation of the ascending aorta into adulthood^27^. In our model, homozygous *Tgfb2* deletion confined to SMCs caused rapid and severe aortic enlargement beginning by 7 weeks, with progressive dilation thereafter. The phenotype was particularly aggressive in the proximal descending aorta, where intramural hematoma and rupture were common, paralleling reports of human *TGFB2* mutations associated with aortic aneurysm, dissection, arterial tortuosity, and cerebrovascular disease^28^.

Loss of *Tgfb2* in postnatal SMCs also produced medial degeneration with elastin fragmentation, SMC dropout, and excessive extracellular matrix (ECM) deposition. Similar histopathology has been described in aortas from patients with *TGFB2* mutations as well as in Marfan and Loeys-Dietz syndromes and genetic and experimental mouse models of thoracic aortopathy ^32^ ^3^ ^2, 9, 27, 40,41^. The pathogenesis of these lesions likely reflects multiple processes. Elastin fragmentation and proteoglycans accumulation compromise structural integrity, while SMC death reduces contractile support ^42–44^.

Dysregulated proteolysis may further contribute to aneurysm progression. Matrix metalloproteinases (MMP2, MMP9, and MMP12), normally produced by SMCs during physiological remodeling, are upregulated during vascular injury by both SMCs and infiltrating immune cells. Imbalance between MMPs and their inhibitors (TIMPs) has been linked to aortic degeneration ^45–47^. Interestingly, in our study, MMP2 and MMP9 activities were not significantly altered in *Tgfb2*-deficient aortas at an early stage of aneurysm formation, suggesting that medial degradation was driven by alternative or parallel mechanisms. TGFβ2 is known to regulate ECM synthesis and adhesion molecules, and to stimulate integrin signaling. This is consistent with recent findings suggesting that short-term complete loss of both TGFβR1/R2 in SMCs of adult mice predisposes to aortic dissection induced by high blood pressure within 7 days of constant infusion and even within 30 minutes of boluses of angiotensin II or norepinephrine^43^. Thus, these data imply that TGFβ2 is a major TGFβ ligand produced by SMCs which plays a continuous role in postnatal vascular homeostasis. Its absence likely impairs elastogenesis and ECM maintenance, facilitating aneurysm formation.

Collagen remodeling was particularly notable in *Tgfb2*cKO mice, with marked accumulation in both media and adventitia. Adventitial thickening is tracked closely with the severity of medial disruption. Increased collagens have been documented in human thoracic aneurysm and dissection specimens of LDS^27, 48, 49^. While collagen deposition may represent a compensatory response to lost wall strength, it carries trade-offs: excess collagen promotes stiffness and predisposes to rupture, while insufficient or poorly cross-linked collagen weakens the wall, both contributing to aneurysm progression.

We also identified striking proteoglycans accumulation in the medial layer of *Tgfb2*cKO mice. Whereas wild-type aortas exhibited uniform basal proteoglycan staining, *Tgfb2*-deficient aortas displayed abnormal accumulation in damaged and dissected regions. These findings mirror observations in Marfan and Loeys-Dietz syndrome and other aneurysmal patients and mouse models, where excess aggrecan and versican have been linked to impaired turnover and tissue swelling ^44^. Physiological proteoglycans interact with ECM components to retain water and buffer hemodynamic forces^50^. However, computational models suggest that abnormal accumulation can elevate interlamellar pressure, disrupt SMC mechanosensing, and compromise wall integrity, thereby accelerating aneurysm development^51^. Collectively, TGFβ2 is required for aortic homeostasis by maintaining healthy proteoglycans levels.

At the molecular level, *Tgfb2* deletion produced broad changes in TGFβ signaling components, SMC contractile markers, and ECM-related genes. Within one week, *Tgfb2* and *Tgfb3* expression were significantly downregulated, while *Tgfb1* and Tgfbr2 were upregulated, with a modest increase in Tgfbr1. In situ hybridization confirmed loss of *Tgfb2* expression in the medial layer. These changes likely reflect compensatory activation of alternative TGFβ signaling components. A similar reparative or compensatory response was reported after postnatal Tgfbr2 deletion in SMCs, which upregulated *Tgfb1*, *Tgfb2*, *Tgfb3*, Tgfbr1, and Tgfbr3^40^. In addition, *Tgfb2*- deficient aortas showed marked downregulation of SMC contractile genes including Myh11, Acta2, Eln, and Lox. Previous studies demonstrated that TGFβ2 promotes SMC gene expression in fibroblasts and myofibroblasts^52^, and drives elongation and contractile protein induction in SMCs and embryonic progenitors^53^. In contrast, exogenous TGFβ1 fails to induce contractile markers in SMCs lacking Tgfbr2^41^, underscoring the essential role of TGFβ2-dependent signaling in maintaining the differentiated, contractile phenotype of vascular SMCs.

All three TGFβ ligands are expressed in the postnatal aorta and signal through the shared receptors TGFβR1 and TGFβR2, as well as common downstream effectors such as SMAD2 and SMAD3. Genetic mutations in *TGFB2* and *TGFB3* have been associated with TAA in Loeys– Dietz syndrome (LDS), whereas no mutations in *TGFB1* have been linked to the aortic aneurysm to date^26^. Most conditional knockout studies in SMCs have focused on receptors (e.g., TGFβR1, TGFβR2) or downstream mediators (e.g., SMAD2, SMAD3), rather than the TGFβ ligands themselves^40, 41, 54–56^. This emphasis reflects both the central role of receptors and Smads as convergence points in TGFβ signaling—whose deletion in SMCs abolishes nearly all TGFβ inputs and produces robust aortic phenotypes—and the added complexity of studying individual ligands due to isoform redundancy and the intricacies of ligand secretion and activation^57–59^.

Dysregulated expression of TGFβ ligands has been reported in TAA patients^60^. Recent in vitro work has shown that different TGFβ isoforms exert broadly similar effects on hiPSC-derived SMC differentiation, yet *TGFB2* uniquely requires TGFβR3 to activate downstream signaling in this context^61^. Our current investigation provides an important first step toward delineating tissue- and isoform-specific functions of the three TGFβ ligands in aortic SMCs. The findings establish a cell-autonomous, postnatal requirement for SMC-derived TGFβ2 in promoting SMC differentiation and maturation, as well as maintaining aortic wall homeostasis.

One of the limitations of this study is the use of *Myh11CreER*^T2^ mice, in which the Cre transgene is inserted on the Y chromosome. As a result, SMC-specific deletion of *Tgfb2* can only be assessed in male mice, restricting the investigation of TGFβ2 function to a single sex. Future studies are needed to determine the role of SMC-derived TGFβ2 in thoracic aortopathy in both sexes. Notably, autosomal *Myh11CreER*^T2^ RAD mice have recently been developed and could be used to overcome this limitation by enabling SMC-specific recombination in both male and female mice^62^. Another limitation is the timing of *Tgfb2* deletion, which in this study was induced at an early age (4 weeks-old). It has been reported that TGF-β neutralization increased aortic rupture in young (postnatal day 16) *Fbn1*^C1041G/+^ mice, but it only had a minor impact at postnatal day 45^63^. Additionally, SMC-specific *Tgfbr2* deletion at 3, 4, 6, and 9 weeks of age leads to progressively less thoracic aortic aneurysm^40, 41^. Since TGFβ signaling may have age-dependent effects, particularly in the context of aortic remodeling and disease progression, it will be important for future research to explore the role of SMC TGFβ2 in older mice to better reflect adult-onset or progressive aortopathy^57, 63^. *Tgfb2*cKO mice develop aneurysms in both ascending and descending aortas, with intramural dissection and/or rupture at the proximal descending aorta. The role of TGFβ2 in second-heart field and neural crest derived SMCs must also be studied as regional differences of TGFβ2 expression in aorta associated with these cell lineages is implicated in pathogenesis of aortic root aneurysm in mice and humans^61, 64, 65^. Together, the data demonstrate that loss of *Tgfb2* in postnatal SMCs induces a highly penetrant aortic phenotype characterized by medial degeneration, elastin fragmentation, excessive collagen and proteoglycan deposition, and profound changes in gene expression. These features closely resemble human syndromic and nonsyndromic thoracic aortic aneurysms, which share aberrant activation of TGFβ1 and SMAD2 signaling. From a translational perspective, these findings highlight *Tgfb2* as a central regulator of aortic wall integrity after birth. The convergence of elastic fiber loss, maladaptive collagen remodeling, and abnormal proteoglycan accumulation suggests that therapeutic strategies aimed at restoring ECM balance or stabilizing SMC contractile phenotype could slow aneurysm progression in *TGFB2*-related disease. Moreover, compensatory upregulation of alternative TGFβ pathway components raises the possibility that selective modulation of ligand-specific TGFβ signaling—rather than global inhibition—may better preserve aortic homeostasis. Collectively, our results establish *Tgfb2* deficiency as a powerful model for studying thoracic aortic aneurysm pathogenesis and underscores the importance of mechanistically targeted interventions for patients with *TGFB2* mutations.

## Author contributions

M. Gebere and M. Azhar conceptualized and designed the study; M. Gebere, M. Chakrabarti, J. Johnson, A. Azhar, and X. Wang conducted experiments and acquired data; M. Gebere, N. Vyavahare, and M. Azhar analyzed and interpreted data; N. Vyavahare and M. Azhar contributed to funding acquisition; and M. Gebere and A. Azhar wrote and edited the manuscript.

## Data availability

The data presented in this study are available upon request.

## Acknowledgements

Research reported in this publication was supported by the University of South Carolina School of Medicine Instrumentation Resource Core Facility (RRID:SCR_024955) and by the Columbia VA Healthcare System. The content is solely the responsibility of the authors and does not necessarily represent the official views of the National Institutes of Health, Columbia VA Healthcare System, or the USC Vice President of Research Office. The authors thank Ms. Lorain Junor for technical assistance in echocardiography. ChatGPT (OpenAI, https://chat.openai.com) was utilized to develop portions of the text and edit the content in this document. The authors subsequently reviewed all the content for its accuracy and originality. Our partnership with ChatGPT upholds strict privacy standards to safeguard data, including proprietary research. Any information entered through our USC-affiliated ChatGPT account stays within our USC system and will not be shared to train the AI model.

